# The Flemmingsome reveals an ESCRT-to-membrane coupling required for completion of cytokinesis

**DOI:** 10.1101/2020.01.15.907857

**Authors:** Cyril Addi, Adrien Presle, Stéphane Frémont, Frédérique Cuvelier, Murielle Rocancourt, Florine Milin, Sandrine Schmutz, Julia Chamot-Rooke, Thibaut Douché, Magalie Duchateau, Quentin Giai Gianetto, Hervé Ménager, Mariette Matondo, Pascale Zimmermann, Neetu Gupta-Rossi, Arnaud Echard

**Author notes:** Equal contributions (Neetu Gupta-Rossi, Arnaud Echard).

## Abstract

Cytokinesis requires the constriction of ESCRT-III filaments on the side of the midbody, where abscission occurs. After ESCRT recruitment at the midbody, it is not known how the ESCRT-III machinery localizes to the abscission site. To reveal novel actors involved in abscission, we obtained the proteome of intact, post-abscission midbodies (Flemmingsome) and identified 489 proteins enriched in this organelle. Among those proteins, we further characterized a plasma membrane-to-ESCRT module composed of the transmembrane proteoglycan syndecan-4, ALIX and syntenin, a protein that bridges ESCRT-III/ALIX to syndecans. The three proteins were highly recruited first at the midbody then at the abscission site, and their depletion delayed abscission. Mechanistically, direct interactions between ALIX, syntenin and syndecan-4 were essential for proper enrichment of the ESCRT-III machinery at the abscission site, but not at the midbody. We propose that the ESCRT-III machinery must be physically coupled to a membrane protein at the cytokinetic abscission site for efficient scission, revealing novel common requirements in cytokinesis, exosome formation and HIV budding.

---

Cytokinesis leads to the physical separation of the daughter cells and concludes cell division. Final abscission occurs close to the midbody (or Flemming body), a prominent structure that matures at the center of the intercellular bridge connecting the two daughter cells and first described by Walther Flemming in 1891^1-9^. The scission occurs not at the midbody itself, but at the abscission site located at distance on one side of the midbody^10-14^. The first scission is usually followed by a second cleavage on the other side of the midbody, leaving a free midbody remnant (MBR)^8, 12-15^. Then, MBRs are either released or wander, tethered at the cell surface for several hours, before being engulfed and degraded by lysosomes^14, 16-19^.

The ESCRT (Endosomal Sorting Complexes Required for Transport) machinery plays a critical and evolutionarily conserved role in cytokinetic abscission, both in Eukaryotes and in Archea^20-31^. This machinery is composed of several protein complexes (ESCRT-0, I, II and III) and culminates with the polymerization of filaments made of ESCRT-III components that contract in the presence of the ATPase VPS4 and ATP^31-35^. Remarkably, ESCRT-III-dependent helices of 17 nm filaments are observed at the abscission site by electron microscopy (EM), and ESCRT-III helical structures are often visible extending from the midbody to the abscission site^10, 36^. Therefore, as in other topologically equivalent ESCRT-III-mediated events, including exosome biogenesis in multivesicular bodies (MVBs), retroviral budding or membrane repair, constriction of ESCRT-III filaments likely drives the final membrane scission during cytokinetic abscission^2-7, 9^.

The midbody plays a fundamental role in cytokinesis, as it constitutes a protein-rich platform that recruits key components for abscission, including the ESCRT machinery^2, 3, 6, 9^. It is well established that the MKLP1 kinesin targets CEP55 to the midbody, which in turn recruits, through both ESCRT-I TSG101 and ESCRT-associated protein ALIX, the entire ESCRT machinery^29, 37^. After this initial recruitment to the midbody itself and prior to abscission, the ESCRT-III machinery is progressively enriched to the future abscission site on the midbody side^12, 13, 20, 21, 25, 30, 31, 36, 38, 39^.

Since MKLP1 and CEP55 are only present at the midbody^38^, it remains elusive how, mechanistically, ESCRT-III components can localize to the abscission site. Another crucial related issue is to reveal how the ESCRT-III filaments could be coupled to the plasma membrane, as final membrane constriction should require their tight association.

Here, we first set up a novel method, using FACS, for purifying intact post-cytokinetic MBRs, and identified by proteomics 489 proteins enriched in this organelle. Among them, we focused on the transmembrane protein syndecan-4 and associated proteins syntenin-1 (hereafter “syntenin”) and ALIX, all highly enriched. Indeed, ALIX directly interacts with ESCRT-III^40^ and we previously showed that syntenin can bind directly and simultaneously to the cytoplasmic tail of syndecans *in vitro*^41, 42^. We thus hypothesized that ALIX-syntenin could mechanistically bridge the ESCRT machinery to the plasma membrane through the transmembrane proteoglycan syndecan-4. Interestingly, overexpression of syndecan-4 mutants that cannot be properly phosphorylated on the cytoplasmic tail was reported to perturb cytokinesis^43^. However, the underlying mechanism is unknown and whether syndecan-4 is actually required for cytokinesis has not been addressed. We here reveal that, together with ALIX, both syndecan-4 and syntenin are required for successful abscission, as they enable the stable recruitment of the ESCRT-III machinery specifically at the abscission site.

## RESULTS

### The Flemmingsome, or proteome of MBRs, reveals new candidates for abscission

The proteome of intercellular bridges from CHO cells previously proved to be a particularly successful approach to identify new proteins required for cytokinesis^44^. However, the use of detergents during the purification steps precluded the recovery of crucial proteins for cytokinesis, for instance the ESCRT components^44^. In order to purify intact midbodies without detergent treatment and thus reveal the complete proteome of these abscission platforms, we took advantage of the fact that released MBRs can be easily detached from the cell surface by EDTA treatment, as we previously reported^14^. Differential centrifugations helped to enrich intact MBRs from EDTA-treated HeLa cells expressing the midbody-localized kinesin GFP-MKLP2^45^ (“MidBody Enriched fraction” or “MBE”, Fig. 1a). To obtain the purest possible MBRs, we developed an original protocol for isolating fluorescent GFP-positive MBRs (“MB+”) from whole cells using FACS sorting (Fig. 1a-b and Supplementary Fig. 1a). In parallel, we isolated small particles of the same size (1-3 μm) and granularity (SSC) but negative for GFP-MKLP2 (“MB-”) (Fig. 1a-b and Supplementary Fig. 1a). Western blot analysis demonstrated that the MB+ population contained highly enriched known midbody proteins [MKLP1, CRIK, PRC1, PLK1, CEP55] and showed no significant contamination, as compared to MB-, total cell lysate (Tot) and MBE fractions, with intracellular compartments [Calreticulin (Endoplamic Reticulum), GM130 (Golgi), Tom22 (mitochondria), HistoneH3 (nucleus), EEA1 (endosomes)] (Fig. 1c and Supplementary Fig. 1b). As expected, proteins such as ALIX, which participates both in cytokinesis and endosomal sorting in interphase were less strikingly enriched (2 fold). Remarkably, cell membrane labeling with Cell Mask and scanning electron microscopy^46^ further demonstrated that MB+ fractions contained membrane-sealed, intact MBRs (Fig. 1d), with very similar shape and length, as observed *in vivo*^14^. Immunofluorescence revealed that the 1-3 μm-sized objects sorted in MB+ were indeed all GFP-positive MBRs and that the respective localization of key cytokinetic proteins [MKLP2, AuroraB, MKLP1, CEP55, RacGAP1, CHMP4B, ALIX, PRC1 and CRIK] was preserved (Fig. 1d). Thus, this novel method to purify MBRs using FACS sorting allowed us to obtain intact and highly pure MBRs, which correspond to midbodies at the time of abscission.

**Figure 1:**
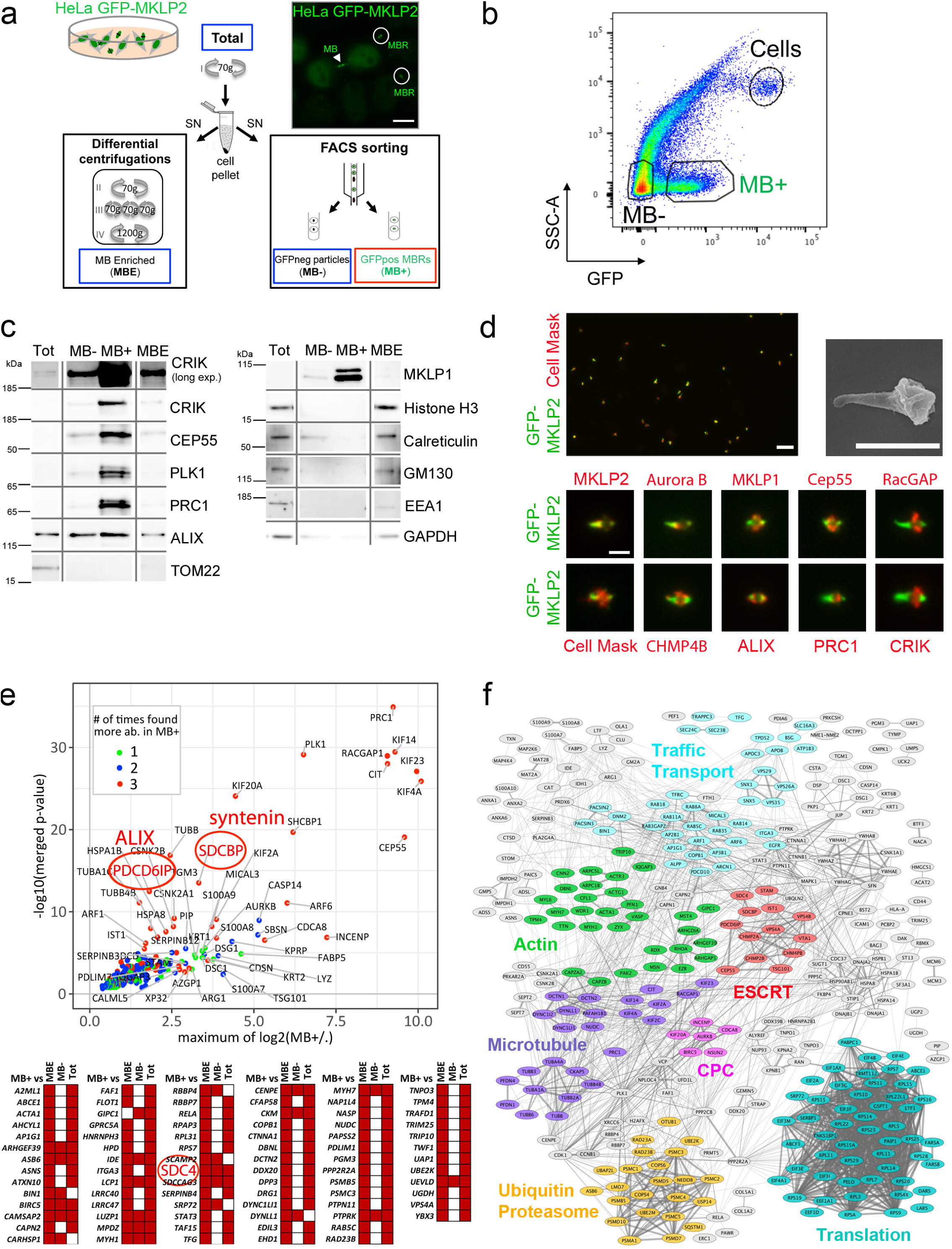
Proteomics of highly pure and intact post-abscission midbodies revealed known and novel proteins enriched in this organelle. (a) Midbody Remnant purification strategy. Starting material was HeLa GFP-MKLP2 cells (upper right picture, GFP signal) that express GFP-MKLP2, a kinesin enriched in intercellular bridges (MB) and midbody remnants (MBRs). Cells were treated with EDTA (Total) to detach MBRs from cells. After centrifugation at 70g to remove most cells, the supernatant (SN) containing MBRs was processed either 1) by differential centrifugations (70g to remove remaining cells, 1200g to pellet MBRs) and led to a MB-enriched fraction (MBE) or 2) subjected to FACS sorting to purify individual GFP-positive MBRs (MB+) and their GFP-negative counterpart (MB-) from SSC-matched particles. (b) Representative pseudo-colored profile of FACS sorting. The GFP-positive particles (MB+; 14% of total) and SSC-matched GFP-negative particles (MB-; 44% of total) were separated from remaining cells (Cells; 1%). See Supplementary Fig. 1a for details. (c) Westernblot analysis of same amounts of protein extracts from total (Tot), midbody enriched (MBE), FACS-sorted MB- and MB+ fractions. Membranes were blotted repeatedly with indicated antibodies. See Supplementary Fig. 1b for un-cropped blots and serial dilutions of the different fractions. For CRIK a long exposure is also displayed. (d) *Upper left panel:* MB+ fraction was analyzed using the membrane marker cell mask. Each object corresponds to an individual midbody positive for both GFP-MKLP2 (green) and cell mask (red) Scale bar: 6 μm. *Upper right panel:* isolated MBR from MB+ fraction observed by scanning electron microscopy. Note that the membrane is intact and sealed on both sides of the midbody. Scale bar: 2 μm. *Lower panels:* immunofluorescence staining of MBRs from MB+ (MKLP2, green) for endogenous proteins or membrane marker (red), as indicated. Scale bar: 2 μm. (e) *Upper graph:* Merged volcano plot of the mass spectrometry analysis showing in x-axis the maximum log2*(fold-change)* measured between MB+ and the other fractions (MBE, MB- or Total) and in y-axis the corresponding –log10*(merged p-value). Lower panel:* Proteins quantitatively present in MB+ but not detected in at least two of the other fractions, and thus not plotted in the merged volcano (See also Supplementary Table 1, TAB4). Colour code upper graph: Proteins significantly enriched when compared with 3 (red), 2 (blue) or 1 (green) of the other fractions (Tot, MBE or MB-samples). ALIX, syntenin (SDCBP or syndecan-binding protein) and syndecan-4 (SDC4) have been highlighted in red circles. (f) Results of the STRING functional association network for the 489 proteins of the *Enriched Flemmingsome* (see Supplementary Table 1, TAB2). Proteins of the *Enriched Flemmingsome* lacking known interactions (152) have not been displayed. Proteins of the same functional category have been colored, as indicated, using Cytoscape. Widths of the edges correspond to the *combined score* of STRING reflecting the confidence that can be placed in each interaction. Only phylogenetic co-occurrence interactions, experimentally determined interactions and database annotated interactions have been considered. CPC: Chromosomal Passenger Complex related.

We next performed proteomic and statistical analysis to 1) identify proteins detected in 7 independent MB+ preparations and 2) identify proteins significantly enriched in these preparations, as compared to MB-, MBE and/or total cell fractions. Since it is notoriously difficult to extract proteins from midbodies^47^, we used SDS to fully solubilize proteins from our different fractions after purification. For mass spectrometry analysis, two complementary methods for sample preparation were used (SDS-PAGE gel/in-gel digestion and eFASP [enhanced Filter-Aided Sample Preparation]/in-solution digestion, Supplementary Fig. 1c). We detected a total of 1732 proteins with at least one unique identified peptide in the MB+ preparations, constituting the *Total Flemmingsome* (Supplementary Table 1, TAB1), a name that we gave as a tribute to W. Flemming.

Among the 1732 proteins in MB+, we defined as the *Enriched Flemmingsome* (Supplementary Table 1, TAB2) a subset of 489 proteins significantly enriched at least 1.3-fold (FDR<5%) as compared to MBE, MB- or Tot (Fig. 1e upper part, Supplementary Fig. 2 and Supplementary Table 1, TAB3) and/or quantitatively present in MB+ but not detected in at least one other fraction (Fig. 1e lower part and Supplementary Table 1, TAB4-5 and Methods). For instance, CRIK was found enriched > 500-fold in MB+ as compared to Total (Supplementary Table 1, TAB2). Interestingly, differential analyses indicated that the most abundant and most significantly enriched proteins, such as MKLP1 (KIF23), MKLP2 (KIF20A), RacGAP1, KIF4A, PRC1, KIF14, PLK1, CEP55, CRIK (CIT) corresponded to well established proteins of cytokinesis (Fig. 1e).

Furthermore, 150 out of the 489 proteins (31%) have been already localized to the furrow, the bridge or the midbody and/or functionally involved in cytokinesis, according to our literature search (Supplementary Table 1, TAB2 and <u>dedicated website</u>). Proteins of the *Enriched Flemmingsome* were highly connected and many fell into known functional categories involved in cytokinesis, such as “actin-related”, “chromosomal passenger complex (CPC)-related”, “microtubule-related”, “traffic/transport-related” or “ESCRT-related” (Fig. 1f and Supplementary Fig. 3). Thus, our approach was highly successful at identifying 150 known cytokinetic proteins and, importantly, revealed 339 new candidates potentially involved in cytokinesis/abscission. In the rest of this study, we decided to focus on a tri-partite complex ALIX (PDCD6IP)-syntenin (SDCBP)-syndecan-4 (SDC4) (Fig. 2a). Indeed, these three proteins were found among the most enriched in MB+ compared to all other fractions (Fig. 1e, highlighted in red).

**Figure 2:**
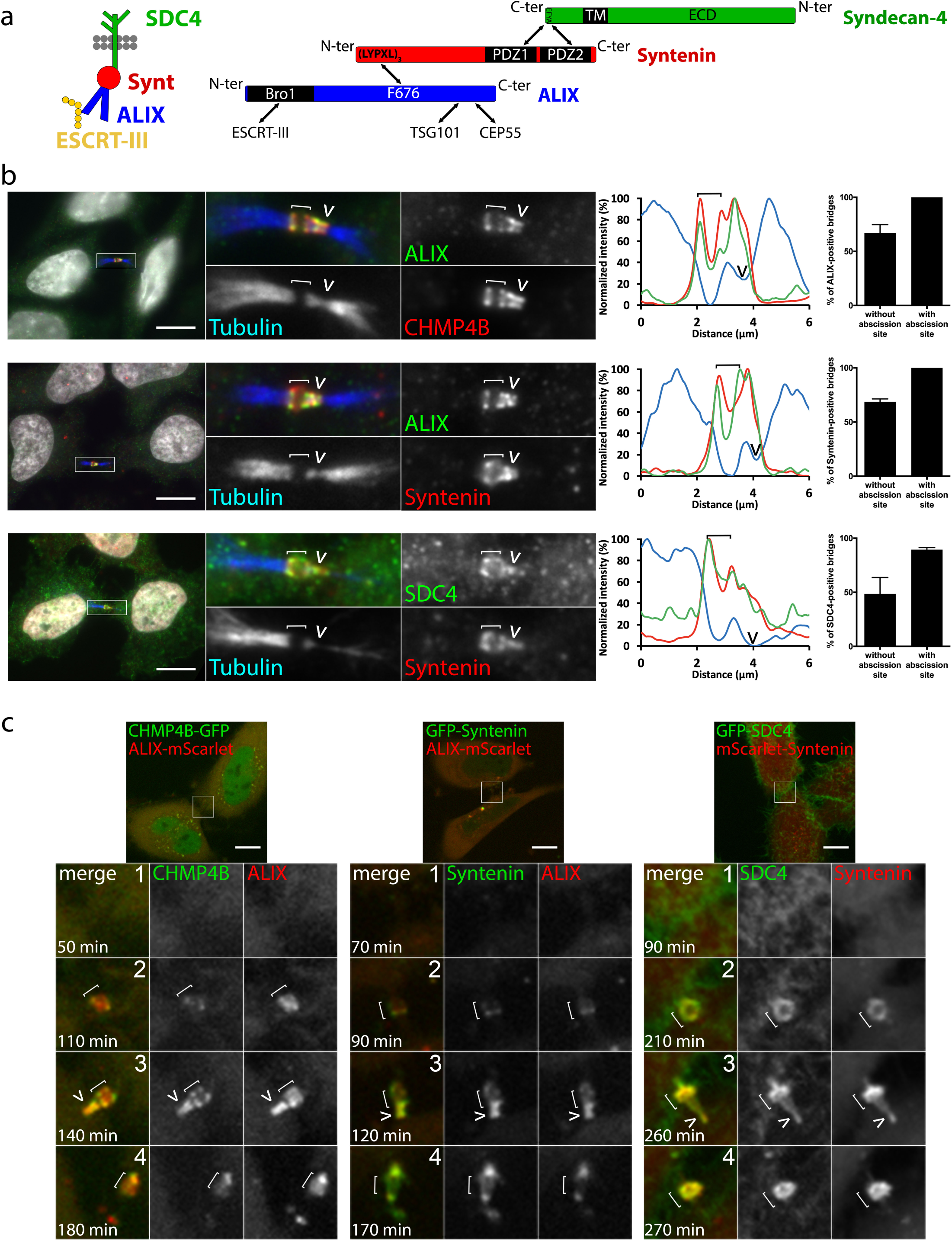
Syndecan-4, syntenin, ALIX and CHMP4B colocalize and are highly enriched first at the midbody then at the abscission site. (a) The ESCRT-III-ALIX-syntenin-syndecan-4 complex. (b) *Left panels:* endogenous localization of ALIX, CHMP4B, syntenin, syndecan-4 (SDC4) and acetyl-tubulin in late bridges displaying abscission site in fixed HeLa cells, as indicated. We used an antibody recognizing the ectodomain of SDC4 for the immunofluorescence. *Middle panels:* Intensity profiles along the bridge of the corresponding images with matched colours from left panels. *Right panels:* Percentage of bridges without and with abscission sites (displaying a pinched tubulin staining on the midbody side) positive for ALIX, syntenin and syndecan-4, as indicated. n ≥ 20 cells, N=3. (c) Snapshots of time-lapse, spinning-disk confocal microscopy movies of cells co-expressing either CHMP4B-GFP/ALIX-mScarlet, GFP-Syntenin/ALIX-mScarlet or GFP-SDC4/mScarlet-Syntenin. Selected time points show cells 1) before the recruitment of the fluorescently-labelled proteins, 2) after their enrichment at the midbody, 3) after their appearance at the abscission site and 4) after abscission. Time 0 corresponds to the time frame preceding furrow ingression. See also corresponding Supplementary videos 1-3. b and c: Scale bars = 10 μm. Brackets and arrowheads mark the midbody and the abscission site, respectively.

### Colocalization of ALIX, syntenin and syndecan-4 along with ESCRT-III at the midbody and at the abscission site

In fixed cells, endogenous ALIX and CHMP4B colocalized as two parallel stripes at the midbody (hereafter figured with white brackets) in bridges without observable abscission sites, as expected (Supplementary Fig. 4a). When bridges mature, the future abscission site also known as secondary ingression site (hereafter pointed with a white arrowhead) appears at the level of pinched and/or interrupted tubulin staining on one side of the midbody. At this late stage, CHMP4B staining extends from the midbody to the abscission site, often appearing as a cone shape structure on one side of the midbody, as previously reported^12, 13, 20, 21, 25, 30, 31, 39^ (Fig. 2b). We found that endogenous ALIX colocalized with CHMP4B both at the midbody and at the abscission site (to our knowledge it is the first time that ALIX is reported at the abscission site) (Fig. 2b). Similarly, we observed colocalization between syndecan-4/syntenin and syntenin/ALIX, both at the midbody and at abscission site (Fig. 2b), using antibodies recognizing endogenous proteins (specific staining confirmed in Supplementary Fig. 4b). Time-lapse spinning-disk confocal microscopy further revealed a striking time-dependent enrichment of these proteins, first at the midbody, then at the abscission site before the cut (Fig. 2c and Supplementary videos 1-3). Thus syndecan-4, syntenin, ALIX and CHMP4B extensively and dynamically colocalized during the terminal steps of cytokinesis, notably at the abscission site.

### Direct and hierarchical recruitment of syndecan-4, syntenin and ALIX to the cytokinetic bridge

We next investigated how syntenin and syndecan-4 are recruited to the intercellular bridge during cytokinesis. First, siRNA-mediated depletion of ALIX strongly reduced the proportion of bridges positive for endogenous syntenin (Fig. 3a-b). In contrast, syntenin depletion had no effect on ALIX recruitment, and syndecan-4 depletion had basically no impact on syntenin or ALIX recruitment (Fig. 3b and Supplementary Fig. 4c). Importantly, upon reintroduction in ALIX-depleted cells, the ALIX F676D mutant that cannot bind to syntenin^41, 48^ was unable to restore the recruitment of endogenous syntenin to the bridge, whereas wild-type ALIX could (Fig. 3c and Supplementary Fig. 4d). Furthermore, a GFP-tagged syntenin ΔALIX (a triple mutant LYP-LAA that cannot interact with ALIX^41^) was no longer recruited to the bridge, whereas GFP-syntenin wild-type and GFP-syntenin ΔSDC (harboring point mutations in PDZ1 and PDZ2 that disrupt the interaction with syndecans^41, 49^) were (Fig. 3d). Thus, ALIX recruits syntenin to the intercellular bridge and this requires a direct interaction between ALIX and syntenin.

**Figure 3:**
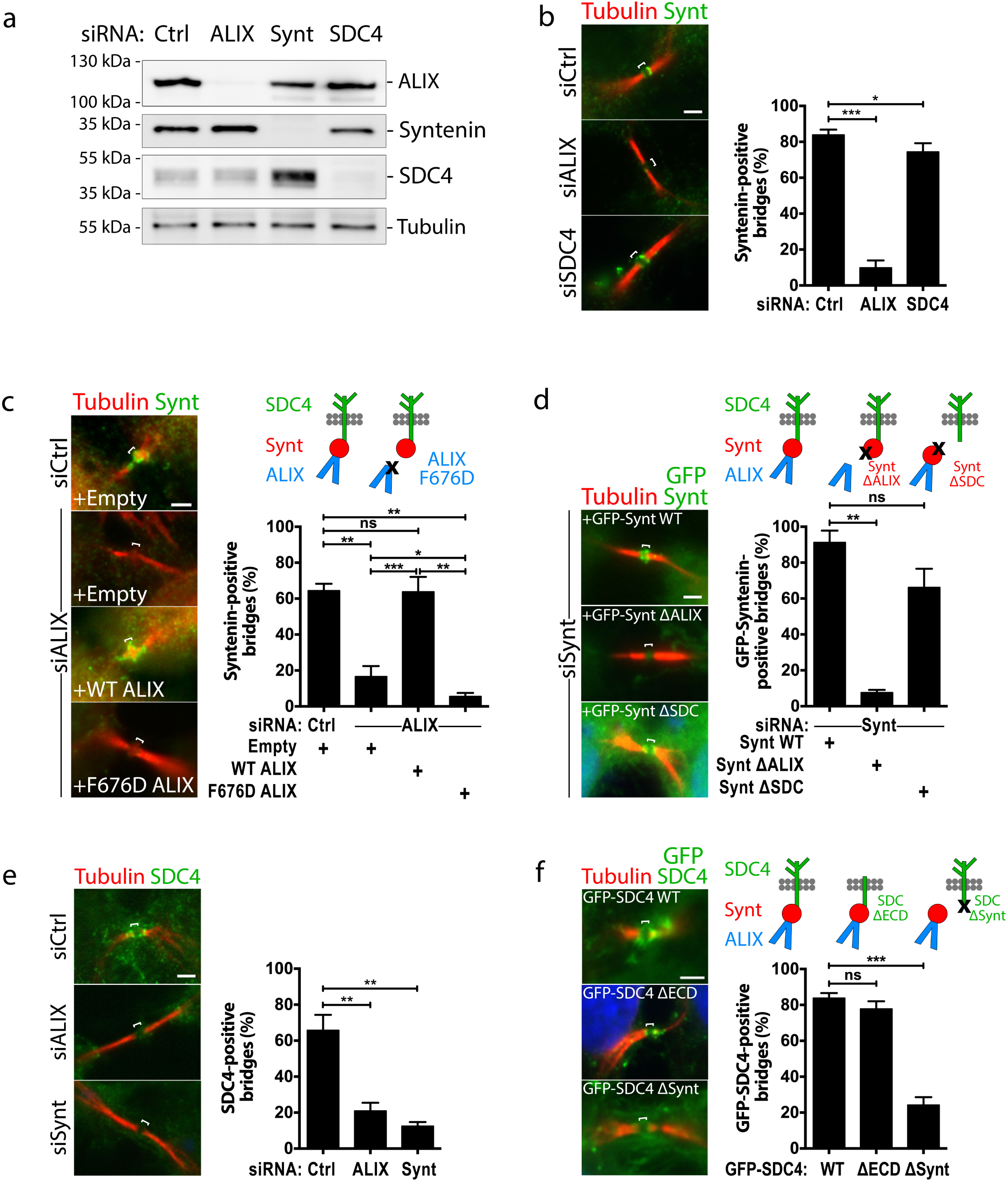
ALIX directly recruits syntenin and syntenin directly recruits syndecan-4 to the cytokinetic bridge. (a) Western blots of cells treated with either control, ALIX, syntenin (Synt) or syndecan-4 (SDC4) siRNAs and revealed with the indicated antibodies. Loading control: β-tubulin. (b) *Left panels:* representative intercellular bridges from control, ALIX or syndecan-4 siRNA-treated cells and stained for acetyl-tubulin and endogenous syntenin, as indicated. *Right panel:* quantification of the % of bridges positive for syntenin after control, ALIX and syndecan-4 depletion. n=31-53 cells, N=3. (c) *Left panels:* representative intercellular bridges from control- or ALIX-depleted cells expressing either control (Empty), wild-type ALIX or ALIX F676D mutant (unable to interact with syntenin), and stained for acetyl-tubulin and endogenous syntenin, as indicated. *Right panel:* quantification of syntenin recruitment in the corresponding conditions. n=25-31 cells, N=3. (d) *Left panels:* representative intercellular bridges from syntenin-depleted cells expressing either GFP-syntenin wild-type, GFP-syntenin ΔALIX (unable to interact with ALIX) or GFP-syntenin ΔSDC (unable to interact with syndecan-4). Acetyl-tubulin and GFP signals are shown. *Right panel:* quantification of GFP-tagged syntenin recruitment in the corresponding conditions. n=14-33 cells, N=4. (e) *Left panels:* representative intercellular bridges from control, ALIX or syntenin siRNA-treated cells and stained for acetyl-tubulin and endogenous syndecan-4, as indicated. *Right panel:* quantification of the % of bridges positive for syndecan-4 after control, ALIX and syntenin depletion. n=30-40 cells, N=3. (f) *Left panels:* representative intercellular bridges from cells expressing either GFP-syndecan-4 wild-type, GFP-syndecan-4 ΔECD (deleted from its entire extracellular domain) or GFP-syndecan-4 ΔSynt (unable to interact with syntenin). Acetyl-tubulin and GFP signals are shown. *Right panel:* quantification of GFP-tagged syndecan-4 recruitment in the corresponding conditions. n=23-41 cells, N=3. b-f: Scale bars = 10 μm. Brackets mark the midbody. ns: non significant, * p < 0.05, ** p < 0.01, *** p < 0.001, Student-t tests.

We next observed that syndecan-4 failed to be correctly recruited at bridges upon ALIX depletion (consistent with the results above) or syntenin depletion (Fig. 3e). Furthermore, GFP-syndecan-4 Δsynt (a mutant deleted of the C-terminal YA motif that is essential for syntenin binding^50^) was no longer recruited to the bridge, while the GFP-syndecan-4 ΔECD (a mutant that lacks the entire extracellular domain) was properly recruited (Fig. 3f). These results indicate that syntenin and its interaction with syndecan-4 are necessary for syndecan-4 localization at the bridge.

Altogether, we conclude that ALIX recruits syntenin that, in turn, recruits syndecan-4 at the intercellular bridge through direct interactions.

### Syndecan-4, syntenin and ALIX are required for efficient cytokinetic abscission

Functionally, time-lapse microscopy demonstrated that abscission was delayed in ALIX-depleted cells (Fig. 4a), as previously reported. Interestingly, abscission was also delayed after either syntenin or syndecan-4 depletions (Fig. 4b-c). Abscission was fully restored by reintroducing either wild-type ALIX, syndecan-4 or syntenin, respectively, ruling out off-target effects (Fig. 4a-c). We conclude that ALIX, syntenin and syndecan-4 are essential for normal abscission.

**Figure 4:**
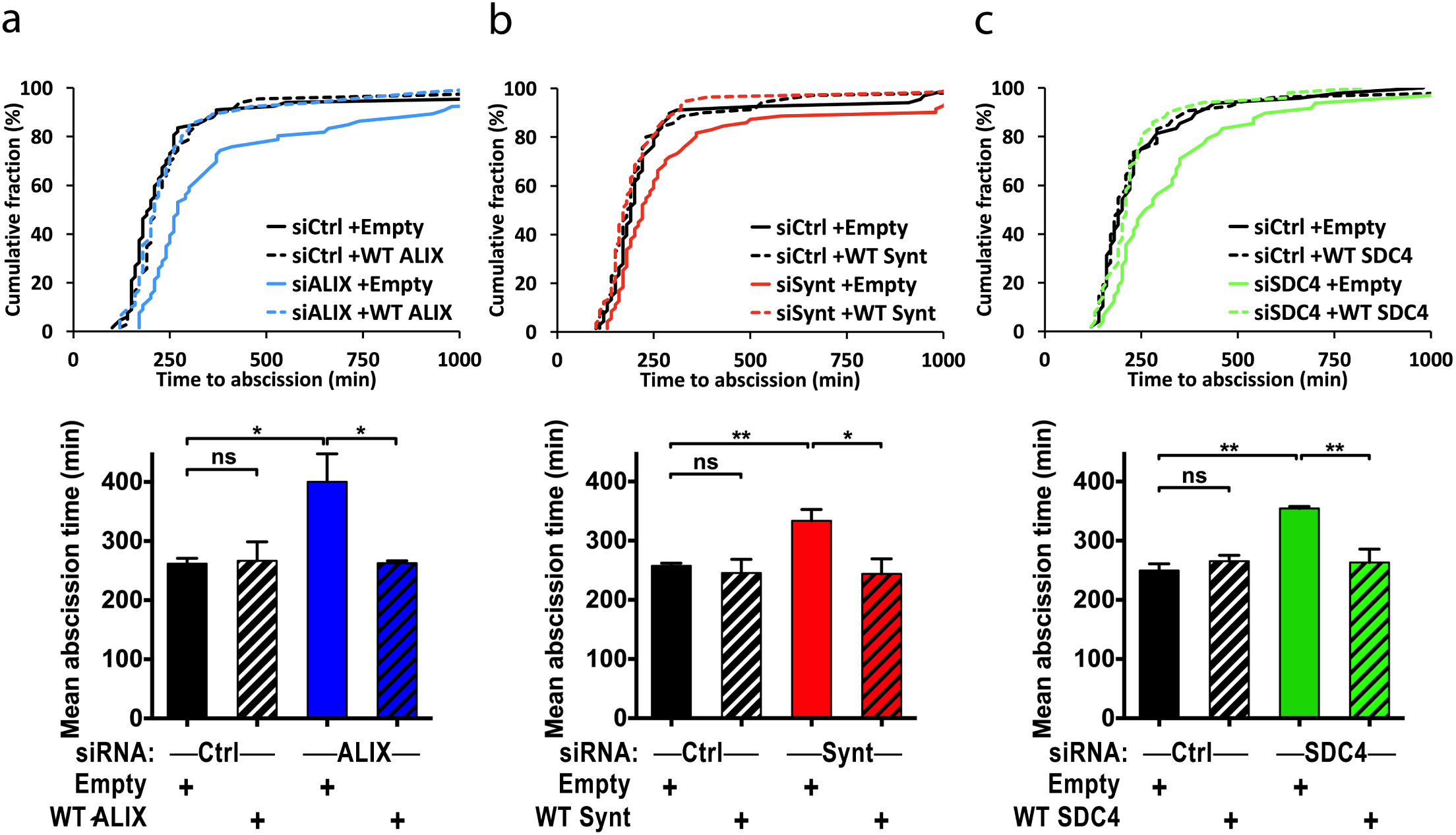
ALIX, syntenin and syndecan-4 are required for successful abscission. (a-c) HeLa cells were treated with control or ALIX (a), syntenin (b) or syndecan-4 (c) siRNAs, then transfected with either empty plasmid or plasmid expressing siRNA resistant transcript encoding the indicated wild-type proteins. Abscission time (from furrow onset to abscission) was determined by phase-contrast time-lapse microscopy. Cumulative plot of the abscission times of a representative experiment (upper panels) and mean abscission times (lower panels) are shown. ALIX depletion: n=159-236 cells, N=3; syntenin depletion: n=180-207 cells, N=3; syndecan-4 depletion: n=130-179 cells, N=3. Time axes were stopped at 1000 min. Comparison of the cumulative abscission time in control vs. depleted cells were significantly different in each condition. p <0.05, using non-parametric and distribution-free Kolmogorov–Smirnov KS tests. Mean abscission times: ns: non significant, * p < 0.05, ** p < 0.01, Student-t tests.

### Depletion of syndecan-4 or syntenin perturbs the recruitment of ESCRT-III at the abscission site but not at the midbody

We then investigated why abscission was delayed after ALIX, syntenin or syndecan-4 depletion. We first quantified the proportion of intercellular bridges with no ESCRT-III at all (early bridges), with ESCRT-III localized only at the midbody (bridges without secondary ingression) and with ESCRT-III both at the midbody and at the abscission site (bridges with constricted/interrupted tubulin staining, as in Fig. 2b). Depletion of either ALIX, syntenin or syndecan-4 considerably reduced the number of bridges with CHMP4B at the midbody + abscission site (Fig. 5a). This is consistent with the observed abscission delay in depleted cells (Fig. 4). Importantly, neither ALIX, syntenin nor syndecan-4 were individually required for correct ESCRT-III recruitment at the midbody itself (the proportion of bridges stuck at this earlier stage was actually increased) (Fig. 5a). As expected, the localization of ESCRT-III at the abscission site was restored in ALIX-, syntenin- or syndecan-4-depleted cells upon expression of siRNA-resistant versions of the corresponding wild-type proteins (Fig. 5b-d). Importantly, the normal localization of ESCRT-III at the abscission site could not be restored by ALIX F676D in ALIX-depleted cells (Fig. 5b), indicating that direct interactions between ALIX and syntenin promote the correct recruitment of ESCRT-III during cytokinesis. Of note, ALIX F676D mutant has been previously reported to rescue the increase in the number of binucleated cells observed after ALIX depletion^21, 22^. In contrast, here, we focused on ESCRT-III recruitment specifically during abscission. Using a second, complementary approach, we found that the syntenin ΔALIX mutant (that cannot bind to ALIX) could not restore normal localization of ESCRT-III during abscission (Fig. 5c), confirming that ALIX-syntenin interaction is indeed important for ESCRT recruitment at the abscission site. Furthermore, defects in ESCRT-III recruitment at the abscission site were also observed with the mutants syntenin ΔSDC (unable to interact with syndecan-4, Fig. 5c) and syndecan-4 Δsynt (unable to interact with and syntenin, Fig. 5d). In contrast, a syndecan mutant lacking the entire ectodomain (but retaining the transmembrane domain + cytoplasmic tail, syndecan-4 ΔECD) localized at the bridge and behaved as wild-type (Fig. 5d). Altogether, we conclude that direct interactions between ALIX and syntenin on one hand, and syntenin and syndecan on the other hand are critical for the recruitment of ESCRT-III specifically at the abscission site but not at the midbody itself.

**Figure 5:**
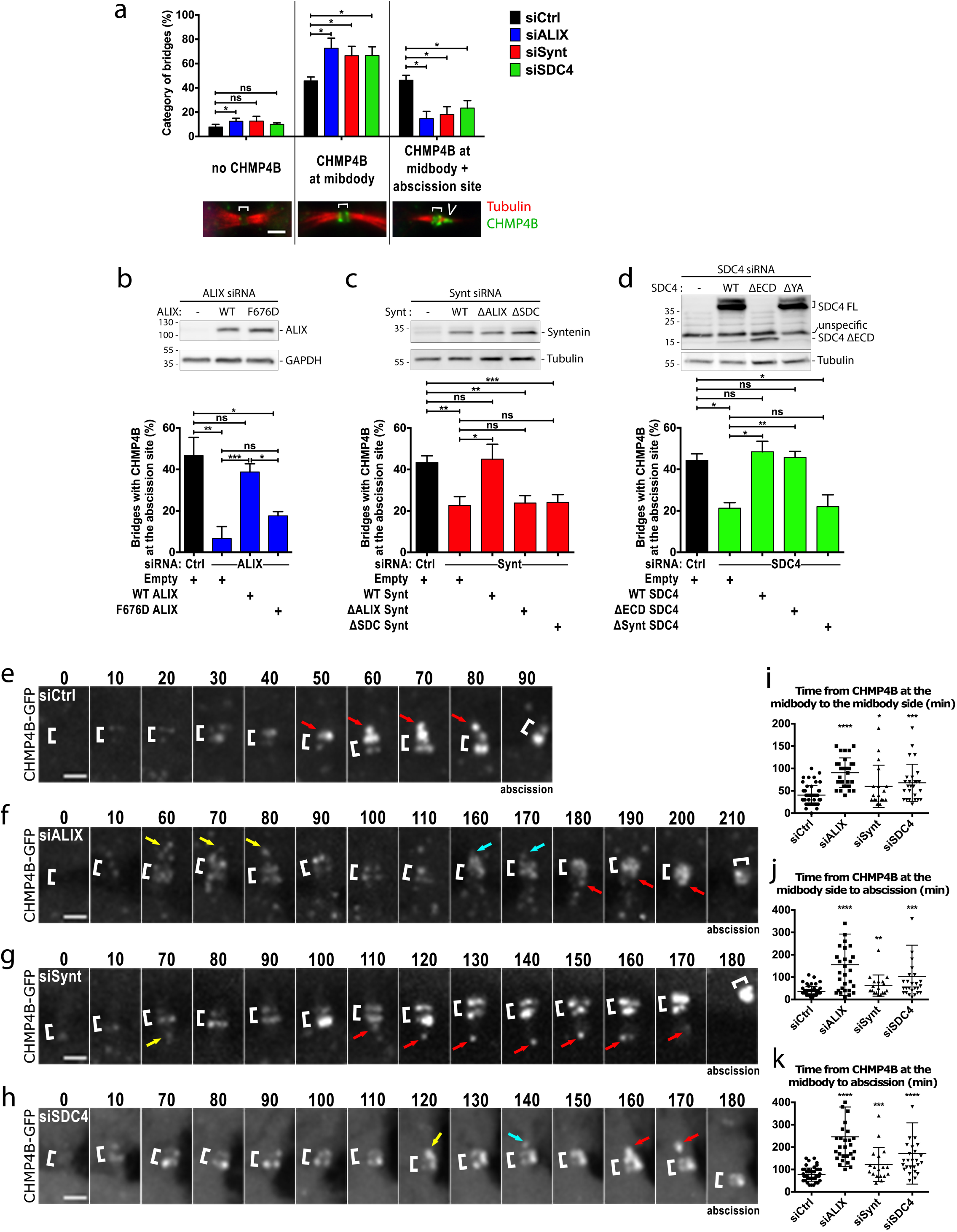
Persistent recruitment of ESCRT-III to the abscission site depends on the syndecan4-syntenin-ALIX module. (a) Cells were treated with either control, ALIX, syntenin or syndecan-4 siRNAs and stained for endogenous CHMP4B and acetyl-tubulin, as indicated. CHMP4B localization in late cytokinetic bridges was classified into three categories: 1) no staining, 2) CHMP4B localized only at the midbody or 3) CHMP4B localized both at the midbody and at the abscission site (representative images are displayed). The proportion of each category was quantified in control and depleted cells, as indicated. (b-d) Cells were depleted for either ALIX (b), syntenin (c) or syndecan-4 (d) and transfected with control plasmid (-) or with plasmids encoding either wild-type or mutant versions of ALIX, syntenin or syndecan-4, as indicated. Upper panels: western blots were revealed with the indicated antibodies for each condition. GAPDH or β-tubulin were used loading controls. Lower panels: quantification of the % of bridges with CHMP4B at the abscission site in each condition, as indicated. n=28-31 cells, N=3 in (b); n=32-103 cells, N=3 in (c); n=33-56 cells, N=3 in (d). (e-h) HeLa cells stably expressing CHMP4B–GFP were treated with either control (e), ALIX (f), syntenin (g) or syndecan-4 (h) siRNAs and recorded by spinning-disc confocal time-lapse microscopy every 10 min. Zooms of the intercellular bridges are displayed. Time 0 corresponds to the frame preceding the arrival of CHMP4B at the midbody. Brackets mark the midbody. Arrows point toward pools of CHMP4B on the midbody side. Red arrows correspond to the CHMP4B leading to abscission (last time frame). Yellow and cyan arrows point to transient and unstable CHMP4B pools observed in depleted cells. See also corresponding Supplementary videos 4-7. (i) Quantification of the time (min) elapsed from CHMP4B enrichment at the midbody to the first appearance of CHMP4B at the midbody side from movies treated as in e-h. Individual dot represents the measurement from a single movie. (j) Quantification of the time elapsed from the first appearance of CHMP4B at the midbody side to the event of abscission from movies treated as in e-h. Individual dot represents the measurement from a single movie. (k) Quantification of the time elapsed from CHMP4B enrichment at the midbody to the event of abscission from movies treated as in e-h. Individual dot represents the measurement from a single movie. a and e-h: Scale bars = 2 μm. ns: non significant, * p < 0.05, ** p < 0.01, *** p < 0.001 Student-t tests. Brackets mark midbodies in a and e-h.

### Syndecan-4-syntenin-ALIX promotes a persistent localization of ESCRT-III at the abscission site

Finally, we investigated why ESCRT-III did not accumulate normally at the abscission site when either ALIX, syntenin or syndecan-4 were depleted, using fluorescent time-lapse microscopy in cells expressing a functional CHMP4B-GFP^31^.

In control cells, CHMP4B-GFP accumulated first at the midbody then appeared on its side as a strong, large, cone-like signal pointing toward the abscission site (Fig. 5e and Supplementary video 4). The appearance of CHMP4B-GFP on the midbody side usually started 40 min after CHMP4B-GFP accumulation at the midbody (Fig. 5i). The CHMP4B-GFP signal at the midbody side remained until abscission, which by mean occurred within 35 min (Fig. 5j). In total, the typical time between CHMP4B-GFP recruitment at the midbody and abscission was 75 min (Fig. 5k). In cells depleted for ALIX (Fig. 5f and Supplementary video 5) the time between the accumulation of CHMP4B at the midbody and abscission largely increased, as compared to control cells (Fig. 5k). Strikingly, during this period, CHMP4B-GFP signal on the midbody side took longer to appear (Fig. 5i). Importantly, we also noticed that the CHMP4B-GFP signal on the midbody side was less prominent and frequently disappeared as if it was unstable, something that we never observed in controls (Fig. 5f, yellow and cyan arrows highlight these transient pools of CHMP4B). As a consequence, the time between the first occurrence of CHMP4B on the midbody side and actual abscission was delayed (Fig. 5j), and abscission eventually occurred without large, cone-shape concentration of CHMP4B at abscission site (Fig. 5f). In syntenin-depleted cells (Fig. 5g and Supplementary video 6) and in syndecan-4-depleted cells (Fig. 5h and Supplementary video 7), quantifications revealed that the times from CHMP4B-GFP recruitment at the midbody to abscission (Fig. 5k), from CHMP4B-GFP recruitment at the midbody to appearance at the midbody side (Fig. 5i) and from first CHMP4B-GFP appearance at the midbody side to abscission (Fig. 5j) were also increased. Furthermore, we again observed an abnormal transient localization of CHMP4B-GFP on the midbody side (Fig. 5g-h yellow and cyan arrows). Although the phenotypes were very similar, all the measured parameters were more affected in ALIX-depleted cells as compared with syntenin- or syndecan-4-depleted cells (Fig. 5i-k), perhaps because ALIX directly recruits ESCRT-III and/or possible alternative pathways that can partially compensate when syntenin or syndecan-4 are depleted. Altogether, we conclude that syndecan-4-syntenin-ALIX promotes ESCRT-III recruitment at the abscission site and is thus key for efficient cytokinetic abscission.

## DISCUSSION

Most proteins involved in cytokinetic abscission strongly accumulate at the midbody^3, 9^. Here, we identified 489 proteins enriched in purified MBRs (*Enriched Flemmingsome*) (Fig. 1, Supplementary Table 1). To isolate MBRs, we developed an original FACS-based protocol that yielded preparations that display three important features. First, the MB+ fractions were highly pure. Second, the MBRs have been obtained from unperturbed cells (no drugs for cell synchronization and no treatment for stabilizing actin or microtubules). Most importantly this purification did not involve any detergents. This allowed us to identify both transmembrane (29 proteins, Supplementary Table 1, TAB2) and membrane associated proteins in this organelle. These three points are key improvements, when comparing with previous proteomes of intercellular bridges, which already proved to be seminal in identifying new essential proteins in cytokinesis^44^. Of note, 68% and 29% of the final list of Skop et al. (160 proteins from CHO cells) were respectively present in our *Total Flemmingsome* (1732 proteins) and *Enriched Flemmingsome* (489 proteins) (Supplementary Table 1, TAB6). The difference in extent of protein recovery between both studies could be explained by 1) differences in cell origins (mouse CHO vs. human HeLa), 2) differences in actual organelles (intercellular bridges before abscission vs. free midbodies as generated at the time of abscission), 3) the membrane integrity of the organelles (use of detergents vs. detergent free, thus preserving cytosolic components), 4) the reduced contaminations and the relative quantification of protein abundance (the *Enriched Flemmingsome* is based on significant enrichment compared to other fractions, including Total cell lysates). We were able to identify many *bona fide* cytosolic (ESCRT-related, actin-related, microtubule-related) and membrane/vesicle associated proteins (e.g. Rab11, Rab35, Rab8, Rab14) involved in cytokinesis^3^ but undetected in previous proteomic analysis (Fig. 1, Supplementary Table 1). Intriguingly, ribosomal proteins were found enriched in the *Flemmingsome*, suggesting a functional interplay between cytokinesis and translation, as anticipated by studies in Drosophila^51, 52^. In addition, proteasome inhibition by MG132 has previously found to delay abscission^53^ and here we revealed the presence of major proteasome components in the *Enriched Flemmingsome* (Fig. 1, Supplementary Table 1).

Importantly, 31% of the proteins present in the *Enriched Flemmingsome* (150 proteins) were already demonstrated to be localized to the furrow, bridge or midbody during cytokinesis and/or to be involved in cytokinesis (Supplementary Table 1, TAB2). This demonstrates the strength of our proteomic study, as well as its potential for identifying new candidates (339) important for cytokinesis. The *Enriched Flemmingsome* thus represents a useful resource for the cytokinesis community and we created a dedicated website (https://flemmingsome.pasteur.cloud/) with continuous updated literature on each hit, as a reference database.

Following up on the *Enriched Flemmingsome*, we found that ALIX, syntenin and syndecan-4 are indeed highly enriched at the midbody, then at the abscission site (Fig. 2). Functional analysis demonstrated that the three proteins are required for proper abscission (Fig. 4), and that they function together for ESCRT-III localization, specifically at the abscission site (Fig. 3). Interestingly, we previously reported that syntenin can directly bridge ALIX to syndecan-1/4 *in vitro*, and that ALIX-syntenin-syndecan is key for budding of intraluminal vesicles in MVBs and exosome production^41^. This likely depends on the ability of ALIX to recruit the ESCRT-III machinery at the neck of intraluminal vesicles in MVBs, but this could not be directly addressed given the small size of these necks. Here, we showed that the same module (ALIX-syntenin-syndecan-4) is reused during cytokinesis at a much larger, micrometric scale, and found that it is actually critical for the stable association of ESCRT-III at the abscission site (Fig. 5).

Our results therefore reveal that ESCRT-III recruitment during cytokinesis relies on two successive, separable phases (Fig. 6). First, MKLP1-associated CEP55 directly interacts with and recruits both TSG101 and ALIX at the midbody^37^ (Fig. 6a). At the midbody, ESCRT-III components are recruited in a redundant manner, directly by ALIX and indirectly by TSG101-ESCRT-II^29^. Accordingly, both TSG101 and ALIX must be simultaneously depleted to prevent ESCRT-III recruitment at the intercellular bridge^29^. However, ESCRT-III localization at the abscission site cannot rely on CEP55, since it is absent from this location (Supplementary Fig. 4e). We now found that ALIX plays a key additional role in cytokinesis: it recruits syntenin, which in turn interacts with the transmembrane protein syndecan-4 (Fig. 6b). Importantly, the ALIX-syntenin-syndecan-4 module is required to stably maintain ESCRT-III components at the abscission site. In the absence of either ALIX, syntenin or syndecan-4, ESCRT-III components can polymerize and extend from the midbody toward the abscission site. However, ESCRT-III recruitment at the abscission site takes longer and is often unstable, which delays abscission (Fig. 5). We thus propose that ALIX-syntenin, by physically coupling ESCRT-III on one hand and the transmembrane protein syndecan-4 on the other hand, helps to maintain ESCRT-III polymers at the abscission site until the final cut (Fig. 6b). Consistently, CHMP4B and syndecan-4 displayed correlated patterns at the abscission site in cells that were fixed shortly before the actual cut (Supplementary Fig. 4f). This chain of interactions appears critical for proper ESCRT-III localization at this site and thus abscission.

**Figure 6:**
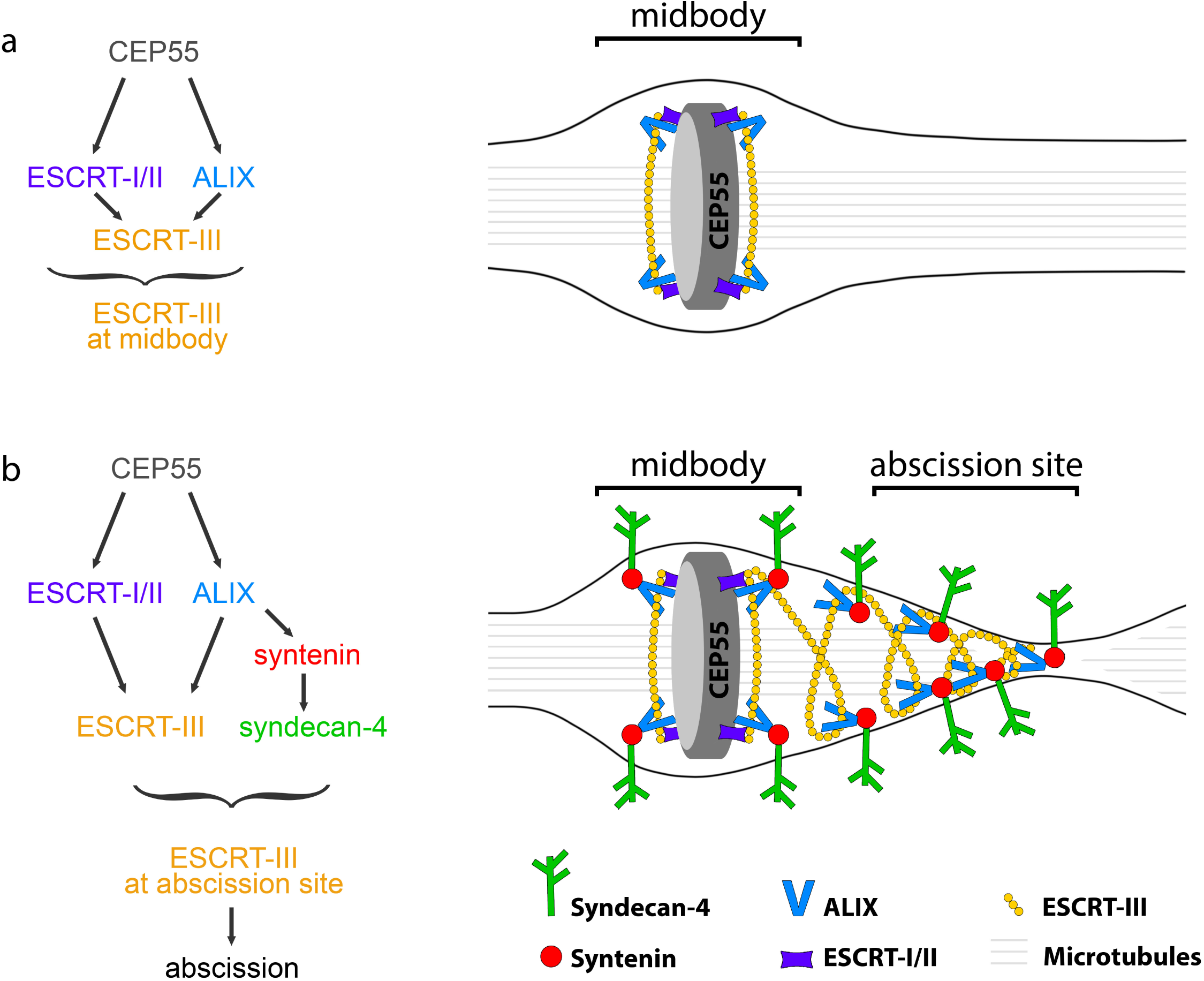
Working model: ALIX-syntenin-syndecan-4 couples the ESCRT-III machinery to the plasma membrane at the abscission site for efficient scission. (a) ESCRT-III localization at the midbody depends on its recruitment by ALIX (blue) and ESCRT-I/-II (violet), which are targeted to the midbody by MKLP1-associated CEP55 (grey). This first step does not require syntenin or syndecan-4. (b) ESCRT-III localization at the abscission site, located on one side of the midbody, depends on the tripartite ALIX-syntenin-syndecan-4 module. ALIX-syntenin (red), by directly coupling ESCRT-III (yellow) on one hand and the transmembrane protein syndecan-4 (green) on the other hand is proposed to help maintain ESCRT-III polymers at the abscission site until the final cut.

In other related ESCRT-dependent membrane scission events, such as exosome formation in MVBs and HIV budding, ALIX also plays a key role^4, 7^. As mentioned above, we found that ALIX-syntenin-syndecan complex is essential for proper exosome scission^41^. Remarkably, HIV appears to have hijacked and simplified this ALIX-syntenin-syndecan module, since the GAG protein (which is tightly associated to the plasma membrane through myristoylation) contains the ALIX-interacting LYpxL motif (that is found 3 times in syntenin), bypassing the need for syndecan-syntenin in the scission step^48, 54-56^. Thus, our study suggests that the coupling of the ESCRT-III machinery to a membrane protein through ALIX-syntenin or equivalent modules represents a fundamental requirement for efficient scission during cytokinesis, exosome formation and retroviral budding.

## ONLINE METHODS

### ell cultures

HeLa cells cl-2 (ATCC) and HeLa GFP-MKLP2^45^ were grown in Dulbecco’s Modified Eagle Medium (DMEM) GlutaMax (31966; Gibco, Invitrogen Life Technologies) supplemented with 10% fetal bovine serum and 1X Penicillin-Streptomycin (Gibco) in 5% CO_2_ condition at 37°C. CHMP4B-GFP and GFP-Syndecan-4 stable cell lines were generated by electroporating HeLa ATCC cells with the corresponding plasmids, followed by G418 selection (Gibco) and selected by FACS sorting.

### Transfections and siRNAs

Plasmids were transfected in HeLa cells for 24 h or 48 h using X-tremeGENE 9 DNA reagent (Roche). For silencing experiments, HeLa cells were transfected once or twice with 25 nM siRNAs for 48h using HiPerFect (Qiagen) following the manufacturer’s instructions. siRNAs against Luciferase (used as control, 5’CGUACGCGGAAUACUUCGA3’), ALIX (5’CCUGGAUAAUGAUGAAGGA3’), Syntenin (5’GAAGGACUCUCAAAUUGCA3’) and Syndecan-4 (5’GUGAGGAUGUGUCCAACAA3’) have been synthetized by Sigma. In rescue experiments, cells were first transfected for 72 h with siRNAs using HiPerFect, then cotransfected by plasmids encoding untagged proteins and either GFP or H2B-GFP to detect transfected cells, using X-tremeGENE 9 DNA reagent for an additional 24 h. SiRNA-resistant versions of ALIX, syntenin and syndecan-4 have been obtained by mutating 6 bp of the siRNA-targeting sequence using NEBaseChanger (NEB).

### Plasmid constructs

Human ALIX, syntenin and syndecan-4 cDNAs were subcloned into Gateway pENTR plasmids and eGFP, mScarlet or untagged transient expression vectors were generated by LR recombination (Thermo Fisher). All point mutations have been generated using NEBaseChanger (NEB), including ALIX F676D, syntenin ΔALIX (Y4A, P5A, Y46A, P47A, Y50A, P51A)^41^, syntenin ΔSDC (K119A, S171H, D172E, K173Q, K203A, K250S, D251H, S252E)^49^ syndecan-4 ΔECD (deleted from E19 to E145 included), syndecan-4 Δsynt (deleted from the last two amino acids, Y197 and A198)^50^. GFP-syndecan-4 was constructed by fusing the eGFP sequence between the syndecan-4 M1-E145 and E142-A198 sequences.

### Western blots

#### Western blot experiments comparing samples used in proteomic studies

Protein extracts from Total, MBE, MB+ and MB-fractions were obtained directly after addition of 2% SDS to the samples. Proteins from approximately 2,5.10^6^ MB+ from FACS were loaded after lysophilisation and resuspension in Laemmli 1x loading buffer on a 4-12% gradient SDS-Page gel (Bio-Rad). Serial dilutions (2-fold) were carried on from 5 to 10 μg extracts from Tot and MBE preparations for comparisons between the samples and SyproRuby protein blot stain (Bio-Rad) used to determine the protein concentrations in the different samples. Lanes with same levels of protein are shown in Fig. 1c and indicated as lane 1 and serial dilutions mentioned from this lane in Supplementary Fig. 1b.

#### Western blot experiments after siRNA treatment were carried out as follows

cells treated with siRNAs were lysed in NP-40 extract buffer (50 mM Tris, pH 8, 150 mM NaCl, 1% NP-40) containing protease inhibitors. 20µg of lysate were migrated in 10% or 4-15% gradient SDS–PAGE gels (Bio-Rad Laboratories), transferred onto PVDF membranes (Millipore) and incubated with corresponding antibodies in 5% milk in 50 mM Tris-HCl pH 8.0, 150 mM NaCl, 0.1% Tween20, followed by HRP-coupled secondary antibodies (1:20,000, Jackson ImmunoResearch) and revealed by chemiluminescence (GE Healthcare). For western blots against syndecan-4, cell extracts were treated with heparinase (AMS.HEP-ENZ III) and chondroitinase (AMS.E1028-02) for 3h at 37°C before migration.

### Immunofluorescence and image acquisition

HeLa cells were grown on coverslips and then fixed either with paraformaldehyde (PFA) 4% for 20 min at room temperature, with methanol for 3 min at −20°C, or with trichloroacetic acid (TCA) 10% for 20 min at room temperature. Cells were then permeabilized with 0.1% Triton-X100, blocked with PBS containing 0.2% BSA and successively incubated for 1h at room temperature with primary (Supplementary Table 2) and secondary antibodies diluted in PBS containing 0.2% BSA. Cells were mounted in Mowiol (Calbiochem). DAPI staining (0.5 mg/ml, Serva). Images were acquired with an inverted TiE Nikon microscope, using a x100 1.4 NA PL-APO objective lens or a x60 1.4 NA PL-APO VC objective lens and MetaMorph software (MDS) driving a CCD camera (Photometrics Coolsnap HQ). Images were then converted into 8-bit images using ImageJ software (NIH). Purified MB+ from FACS were concentrated at 1200g and a 5μl-drop was incubated overnight on a glass coverslip. The MB+ MBRs were processed for immunofluorescence as described above for cells. Cell Mask (C10045, ThermoFisher) staining was performed on the GFP-MKLP2 adherent cells as indicated by the manufacturer, and then the MB+ MBRs purified by FACS as described above.

### Time-lapse microscopy

For time-lapse phase-contrast, HeLa cells were plated on glass bottom 12-well plates (MatTek) and put in an open chamber (Life Imaging) equilibrated in 5% CO2 and maintained at 37°C. Time-lapse sequences were recorded every 10 min for 48 h using a NikonEclipse TiE inverted microscope with a x20 0.45 NA Plan Fluor ELWD controlled by Metamorph software (Universal Imaging). For time-lapse fluorescent microscopy, images were acquired using an inverted Eclipse TiE Nikon microscope equipped with a CSU-X1 spinning disk confocal scanning unit (Yokogawa) and with a EMCCD Camera (Evolve 512 Delta, Photometrics). Images were acquired with a x60 1.4 NA PL-APO VC and MetaMorph software (MDS).

### Statistical analysis

All values are displayed as mean ± S.D. for at least three independent experiments (as indicated in the figure legends). Significance was calculated using unpaired t-tests, as indicated. For abscission times, a non-parametric Kolmogorov– Smirnov test was used. In all statistical tests p>0.05 was considered as not significant. By convention, * p<0.05; ** p<0.01 and *** p<0.001.

### Sample preparation for mass spectrometry

Cells were detached from flasks with 0.05% trypsin diluted in 0.02% EDTA (25300; Gibco, Invitrogen Life Technologies) and plated at 8.10^5^ cells/well on 10-cm dishes for 3 days. Cells were rinsed 3-times with HBSS and then incubated in 2mM EDTA for 10 min at 37°C in order to detach MBRs from the cell surface. The “Total” fraction represented the whole fraction of detached cells including the EDTA-supernatant (SN). The cells were pelleted by centrifugation (5 min at 70 g). The “Midbody-Enriched fraction (MBE)” was adapted from ref.^14^: the supernatant from the first centrifugation was centrifuged again (5 min at 70g) and SN from this step was aliquoted to 300 μl for another centrifugation (10 min at 70g). The MBRs were concentrated from the last supernatant by 60 min centrifugation at 1200 g.

### FACS experimental procedures

For FACS sorting, the supernatant from the first 70g centrifugation was collected. Sorting of MBRs was performed on a BD Biosciences FACS ARIA III. Neutral Density filter 1.0 has been used to detect small particles. MBRs were gated on a pseudo-color plot looking at GFP versus SSC-A parameters, both in log scales. Cells have been excluded from the sorting gates after analysis of an unstained cell suspension as control. The MB+ (GFP-positive population) and MB-(GFP-negative population) populations purified from FACS were concentrated by 60 min centrifugation at 1200 g. The proteins from all the samples were solubilized in 2% SDS and further prepared for in-gel or in-solution digest.

### In gel digestion

In-gel digestion was performed by standard procedures^57^. Proteins (10 μg) were loaded on a SDS-PAGE gel (4-20% gradient, Expedeon). The electrophoretic migration of the gel was stop after the stacking and the gel was stained with Coomassie Blue (InstantBlue™, Expedeon) and each lane was cut into 3 gel bands. Gel slices were washed several times in 50 mM ammonium bicarbonate, acetonitrile (1:1) for 15 min at 37 °C. Disulfide bonds were reduced with 10mM DTT and cysteine alkylated with 55mM IAA. Trypsin (Promega) digestion was performed overnight at 37 °C in 50 mM ammonium bicarbonate. Peptides were extracted from the gel by two incubations in 10% formic acid, acetonitrile (1:1) for 15 min at 37 °C. Extracts were dried in a Speed-Vac, and resuspended in 2% acetonitrile, 0.1% formic acid prior to LC-MS/MS analysis. For each sample (Total, MBE, MB+ and corresponding MB-) 5 independent preparations were run on SDS-PAGE; an experimental replicate was made as an internal control for the MB+/MB-samples (numbered 3 and 4, Supplementary Table 1, TAB3).

### In-solution digestion (eFASP)

Other protein samples extracted in SDS were digested using eFASP protocol^58^. Filter units and collection tubes were incubated overnight in passivation solution: 5% (v/v) TWEEN®-20. All buffer exchanges were carried out by centrifugation at 14 000g for 10 min. Briefly, 10 µg of proteins from each sample was transferred into 30 000 MWCO centrifugal unit (Microcon® Centrifugal Filters, Merck) completed until 200 μL with exchange buffer (8 M urea, 0.2% DCA, 100 mM ammonium bicarbonate pH8.0). Disulfide bonds were reduced with 5mM TCEP (Sigma) for 1h. Proteins were buffer-exchanged with three rounds of 200 μL of exchange buffer. Buffer was then exchange for the alkylation buffer (50 mM iodoacetamide, Urea 8M, 100mM ammonium bicarbonate pH 8) in the dark for 1h. One 200 μL exchange buffer exchanges were used to remove the alkylating agent, followed by three buffer exchanges with 100 μL of digestion buffer (0,2 % DCA / 50 mM ammonium bicarbonate buffer pH 8). 100 μL of digestion buffer containing 1:50 ratio of sequencing-grade modified trypsin (Promega) to amount of protein was added to the retentate. Proteolysis was carried out at 37 °C overnight. Three rounds of 50 μL of recovery buffer (50 mM ammonium bicarbonate pH8.0) was used to elute the peptide-rich solution. Then peptides were process as described in ref.^58^ and resuspended in 2% acetonitrile, 0.1% formic acid prior to LC-MS/MS analysis. For Total and MBE, 3 independent replicates were made; for MB+ and corresponding MB-FACS-sorted samples 2 independent replicates were processed for eFASP (Supplementary Table 1, TAB3).

### Mass spectrometry analysis

Tryptic peptides from in-gel digestion were analyzed on a Q Exactive HF instrument (Thermo Fisher Scientific, Bremen) coupled with an EASY nLC 1000 chromatography system (Thermo Fisher Scientific). Sample was loaded on an in-house packed 50 cm nano-HPLC column (75 μm inner diameter) with C18 resin (1.9 μm particles, 100 Å pore size, Reprosil-Pur Basic C18-HD resin, Dr. Maisch GmbH, Ammerbuch-Entringen, Germany) after an equilibration step in 100 % solvent A (H2O, 0.1 % FA). Peptides were first eluted using a 2 to 7 % gradient of solvent B (ACN, 0.1 % FA) during 5min, then a 7 to 23 % gradient of solvent B during 80 min, a 23 to 45 % gradient of solvent B during 40 min and finally a 45 to 80 % gradient of solvent B during 5 min all at 250 nL.min^-1^ flow rate. The instrument method for the Q Exactive HF was set up in the data dependent acquisition mode. After a survey scan in the Orbitrap (resolution 60 000), the 10 most intense precursor ions were selected for HCD fragmentation with a normalized collision energy set up to 28. Charge state screening was enabled, and precursors with unknown charge state or a charge state of 1, 7, 8 and >8 were excluded. Dynamic exclusion was enabled for 45 s.

Tryptic peptides from eFASP digestion were analyzed on a Q Exactive plus instrument (Thermo Fisher Scientific, Bremen) coupled with an EASY nLC 1000 chromatography system (Thermo Fisher Scientific). Sample was loaded on an in-house packed 50 cm nano-HPLC column (75 μm inner diameter) with C18 resin (1.9 μm particles, 100 Å pore size, Reprosil-Pur Basic C18-HD resin, Dr. Maisch GmbH, Ammerbuch-Entringen, Germany) after an equilibration step in 100 % solvent A (H2O, 0.1 % FA). Peptides were first eluted using a 2 to 5 % gradient of solvent B (ACN, 0.1 % FA) during 5min, then a 5 to 22 % gradient of solvent B during 150 min, a 22 to 45 % gradient of solvent B during 60 min and finally a 45 to 80 % gradient of solvent B during 10 min all at 250 nL.min^-1^ flow rate. The instrument method for the Q Exactive Plus was set up in the data dependent acquisition mode. After a survey scan in the Orbitrap (resolution 70 000), the 10 most intense precursor ions were selected for HCD fragmentation with a normalized collision energy set up to 28. Charge state screening was enabled, and precursors with unknown charge state or a charge state of 1, 7, 8 and >8 were excluded. Dynamic exclusion was enabled for 45 s.

### Data processing for protein identification and quantification

All data were searched using Andromeda^59^ against a Human Uniprot database (downloaded in 20150818, 20204 entries), usual known mass spectrometry contaminants and reversed sequences of all entries. Andromeda searches were performed choosing trypsin as specific enzyme with a maximum number of two missed cleavages. Possible modifications included carbamidomethylation (Cys, fixed), oxidation (Met, variable) and Nter acetylation (variable). The mass tolerance in MS was set to 20 ppm for the first search then 4.5 ppm for the main search and 20 ppm for the MS/MS. Maximum peptide charge was set to seven and five amino acids were required as minimum peptide length. The “match between runs” feature was applied for samples having the same experimental condition with a maximal retention time window of 1 minute. One unique peptide to the protein group was required for the protein identification. A false discovery rate (FDR) cutoff of 1 % was applied at the peptide and protein levels. Quantification was performed using the XIC-based LFQ algorithm with the Fast LFQ mode as described in ref.^60^. Unique and razor peptides, included modified peptides, with at least 2 ratio count were used for quantification.

### Statistical analysis for proteomic studies

For the differential analyses, proteins identified in the reverse and contaminant databases and proteins “only identified by site” were first discarded from the list of identified proteins. Then, proteins exhibiting fewer than 2 quantified values in at least one condition were discarded from the list. After log2 transformation of the leftover proteins, LFQ values were normalized by median centering within conditions (*normalizeD* function of the R package *DAPAR*^61^). Remaining proteins without any LFQ value in one of both conditions have been considered as proteins quantitatively present in a condition and absent in another (Supplementary Table 1, TAB4). They have therefore been set aside and considered as differentially abundant proteins. Next, missing values were imputed using the imp.norm function of the R package norm^62^. Proteins with a fold-change under 1.3 have been considered not significantly differentially abundant. Statistical testing of the remaining proteins (having a fold-change over 1.3) was conducted using a *limma* t-test^63^ thanks to the R package *limma*^64^. An adaptive Benjamini-Hochberg procedure was applied on the resulting p-values thanks to the function *adjust*.*p* of R package *cp4p*^65^ using the robust method described in ref.^66^ to estimate the proportion of true null hypotheses among the set of statistical tests. The proteins associated to an adjusted p-value inferior to a False Discovery Rate (FDR) of 5% have been considered as significantly differentially abundant proteins. Finally, the proteins of interest are therefore those which emerge from this statistical analysis supplemented by those which are considered to be absent from one condition and present in another. Results of these differential analyses are summarized in Supplementary Fig. 2. The merged volcano plot (Fig. 1e) is a summary of the six comparisons MB+ vs control (Neg, MBE or Total; using either eFASP or Gel). The x-axis represents the maximum log2 fold-change between MB+ and the different controls. The “*merged p-value*” (y-axis) has been obtained using the Fisher’s method from the different p-values that have been measured in the comparisons. Note that these two quantities have been computed only when data are available.

### UpsetR graph and functional association network

The Upset graph (Supplementary Fig. 1c) represents the distribution of the significant proteins coming from the different statistical analyzes. The Venn diagram represents the numbers of differentially abundant proteins in function of the kind of experiment (Gel or eFASP). Functional association network was determined by STRING^67^ and displayed using Cytoscape^68^.

## ACKNOWLEDGMENTS

We thank R. Basto, G. Hickson, J. Mathieu, J-R Huyhn, R. Shaughnessy and T. Wai for critical reading of the manuscript; the Echard Lab members for helpful discussions; the Recombinant antibodies platform (TAb-IP, Institut Curie, Paris) and the DHSB (University of Iowa) for antibodies. GFP-MKLP2 cells were from the Hyman Lab MPI-MCBG Dresden^45^. We thank the imaging facilities Imagopole and Ultrapole, Institut Pasteur. This work has been supported by Institut Pasteur, CNRS, and ANR (AbCyStem, Cytosign) to A.E. C.A. and A.P. received a fellowship from the Doctoral School Complexité du Vivant ED515, contrats n°2611 bis/2016 (A.P.), 2412/2016 and AMX (C.A.).

## AUTHOR CONTRIBUTIONS

C.A. carried out and analyzed the experiments presented in Fig. 2, 3, 4, 5 and Supplementary Fig. 4; N.GR. and A.P. in Fig. 1a-d and Supplementary Fig. 1a-b; S.F. in Fig. 1d; F.M. in Fig. 3c; N.GR. and S.S. setup the FACS-purification protocol for Fig. 1b, Supplementary Fig. 1a; N.GR and A.P. setup MBE protocols. F.C. and M.R. assisted with technical help. T.D, M.D. and M.M. carried out the mass spectrometry studies; Q.G.G. did statistical analyses and Fig. 1e-f, Supplementary Fig. 1c, 2 and 3; N.GR., Q.G.G., M.M., M.D., T.D. contributed to the Supplementary Table 1; H.M. created the dedicated website. We acknowledge the help of Thomas Menard from the IT Department at the Institut Pasteur for this work. P.Z. provided reagents and helpful discussions. A.E. conceived the project and secured funding. N.GR. and A.E. supervised the proteomic data; A.E. supervised the other data. A.E. wrote the manuscript with the help of C.A, N.GR., M.D., T.D., Q.G.G., M.M. and P.Z.

## COMPETING INTERESTS

Authors declare no competing interests.

## DATA AND MATERIAL AVAILABILITY

The mass spectrometry proteomics data have been deposited to the ProteomeXchange Consortium via the PRIDE^69^ partner repository with the dataset identifier PXD013219. The Flemmingsome Website: https://flemmingsome.pasteur.cloud/

## FIGURE LEGENDS

**Supplementary Fig. 1:**
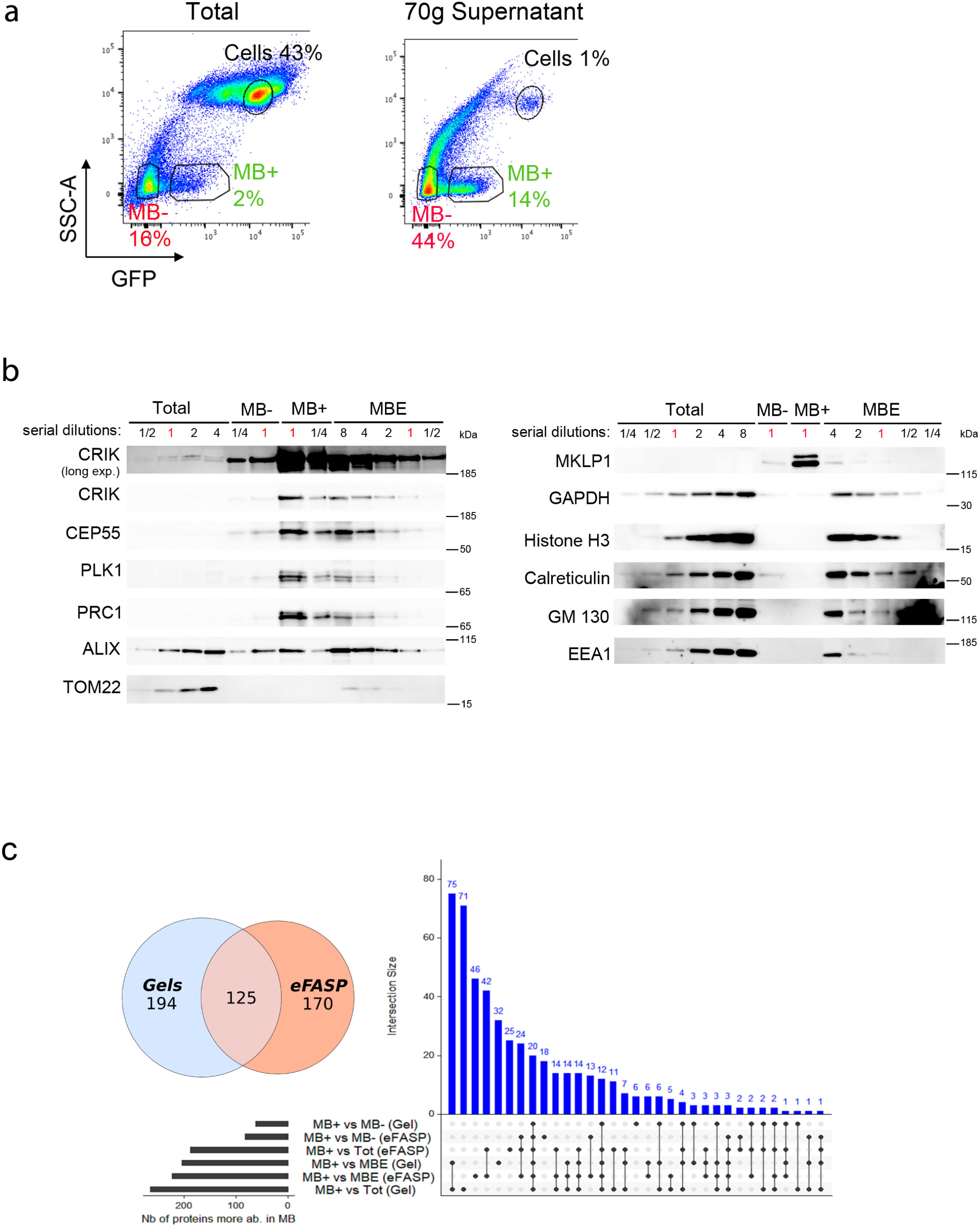
Isolation and characterization of MBRs from GFP-MKLP2 cells. (a) Isolation of MBRs by Flow Cytometry. *Left panel*. GFP-MKLP2 cells were treated with EDTA (to detach cells from the substrate and MBRs from cell surface) (Total) and analyzed by FACS (see also Fig. 1a). Gates show 43% Cells and distinct populations of small GFP-positive MBRs (2% MB+) and GFP-negative particles (16% MB-) in the low SSC population. *Right panel*. For comparison: supernatant from previous condition after a 70g centrifugation step, leading to cell depletion (1% remaining cells) and enrichment of MB+ (14%) and MB-(44%) populations (same graph as in Fig. 1b). The 70g supernatant was used for the sorting of MB+ and MB-fractions for the proteomics analysis. Several FACS independent experiments were pooled and MB+ and MB-particles concentrated at 1200g to obtain enough MB+ and MB-protein extracts for the proteomics analysis. (b) Uncropped Western blots of Fig. 1c, with indicated serial dilutions of protein extracts from total (Tot), midbody enriched (MBE), FACS-sorted MB- and MB+ fractions. The same membrane was blotted repeatedly with indicated antibodies. Sypro-Ruby staining was used to calibrate the protein quantity before membrane blotting with the indicated antibodies (same amount of proteins in lanes labeled as “1”). (c) UpSet plot and Venn diagram showing the significantly more abundant proteins in the MB+ than in the 3 different used controls (MB-, MBE, Total) with 2 different sample preparation techniques (eFASP or Gels). When merging all the results, 489 proteins were found significantly enriched in at least one of the comparisons: 194 are found uniquely enriched using Gels and 170 uniquely when using eFASP while 125 are found enriched with both techniques.

**Supplementary Fig. 2:**
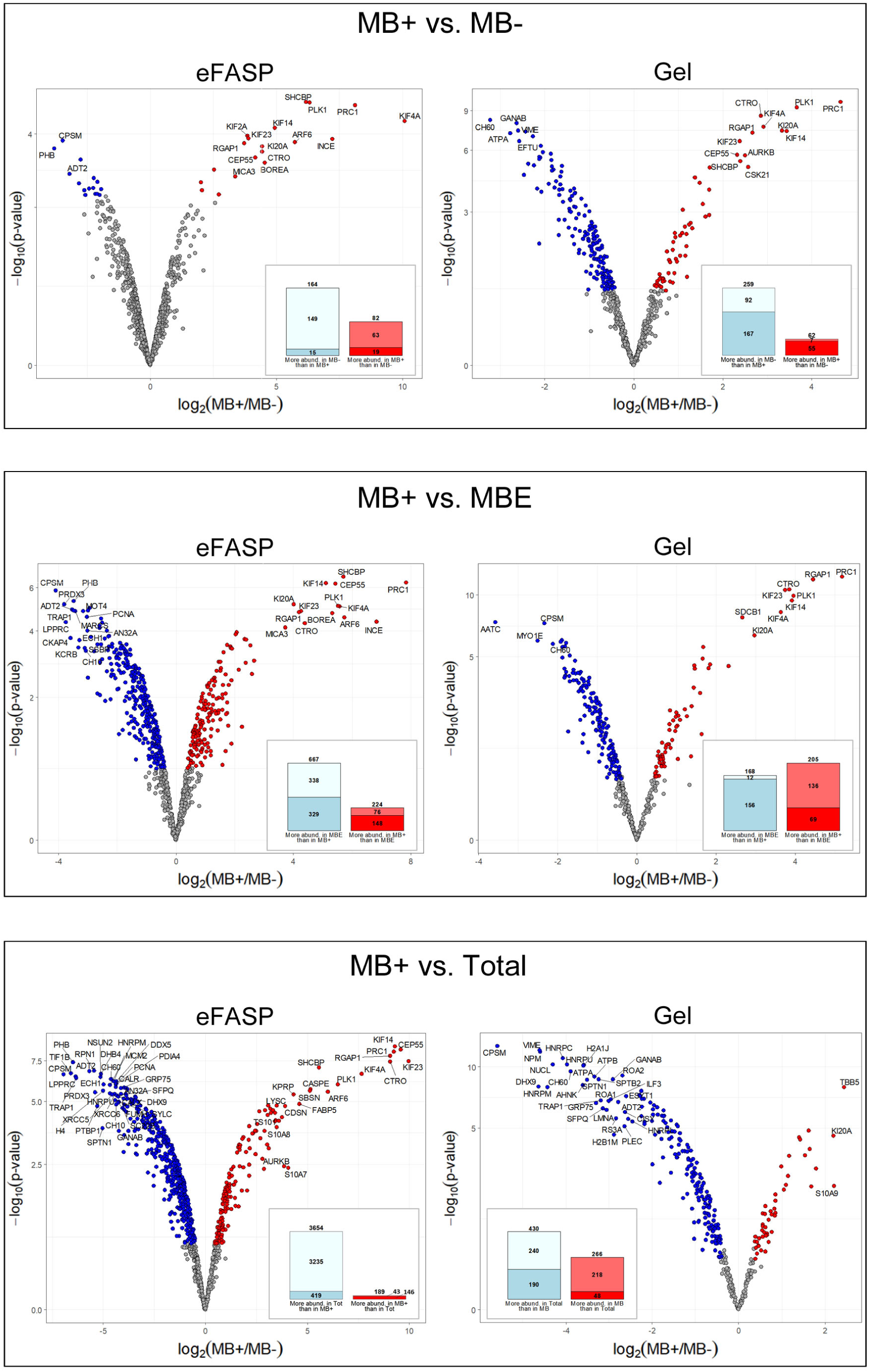
Identification of the *Enriched Flemmingsome* using comparisons between proteomes of MB+ vs. other fractions. Volcano plots showing the proteins that are significantly more abundant in MB+ or in the different used controls [MB-, MBE or Total and using either eFASP or Gels] and quantified in both conditions. The proteins that are significantly more abundant in MB+ are represented in red and the ones significantly more abundant in controls are in blue. The x-axis represent the log2 ratio between average intensities in MB+ and average intensities in controls and the y-axis is the −log10(p-value) where the p-value has been computed with a LIMMA t-test. Bar plots represent the total number of proteins that are more abundant in MB+ than in the different controls (proteins of volcano plot + proteins only quantified in a condition) and vice versa (lightblue: proteins only quantified in the control, blue: proteins significantly more abundant in the control and selected by the statistical test, lightred: proteins only quantified in MB+, red: proteins significantly more abundant in MB+ and selected by the statistical test).

**Supplementary Fig. 3:**
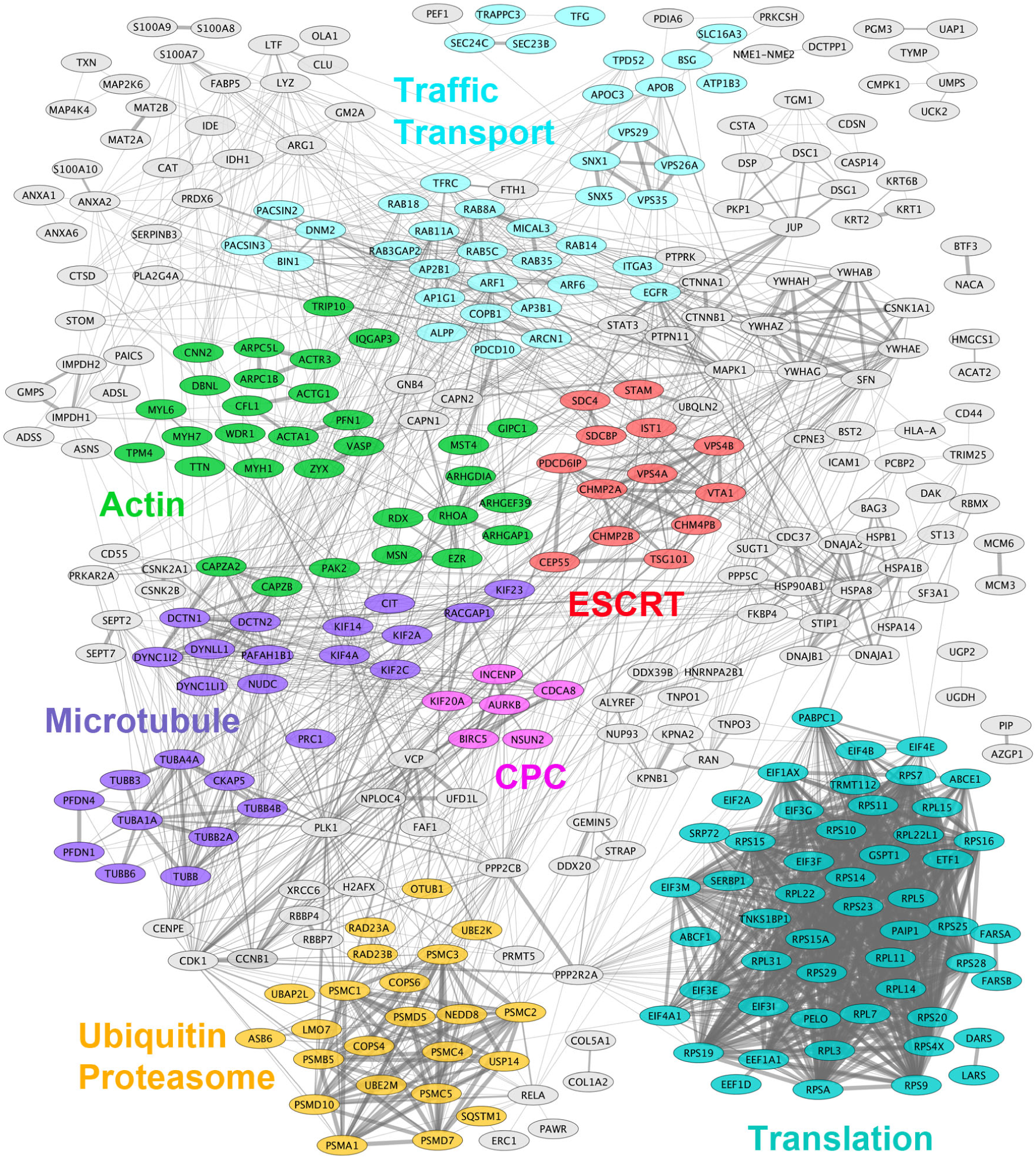
Zoom on the association network displayed in Fig. 1f.

**Supplementary Fig. 4:**
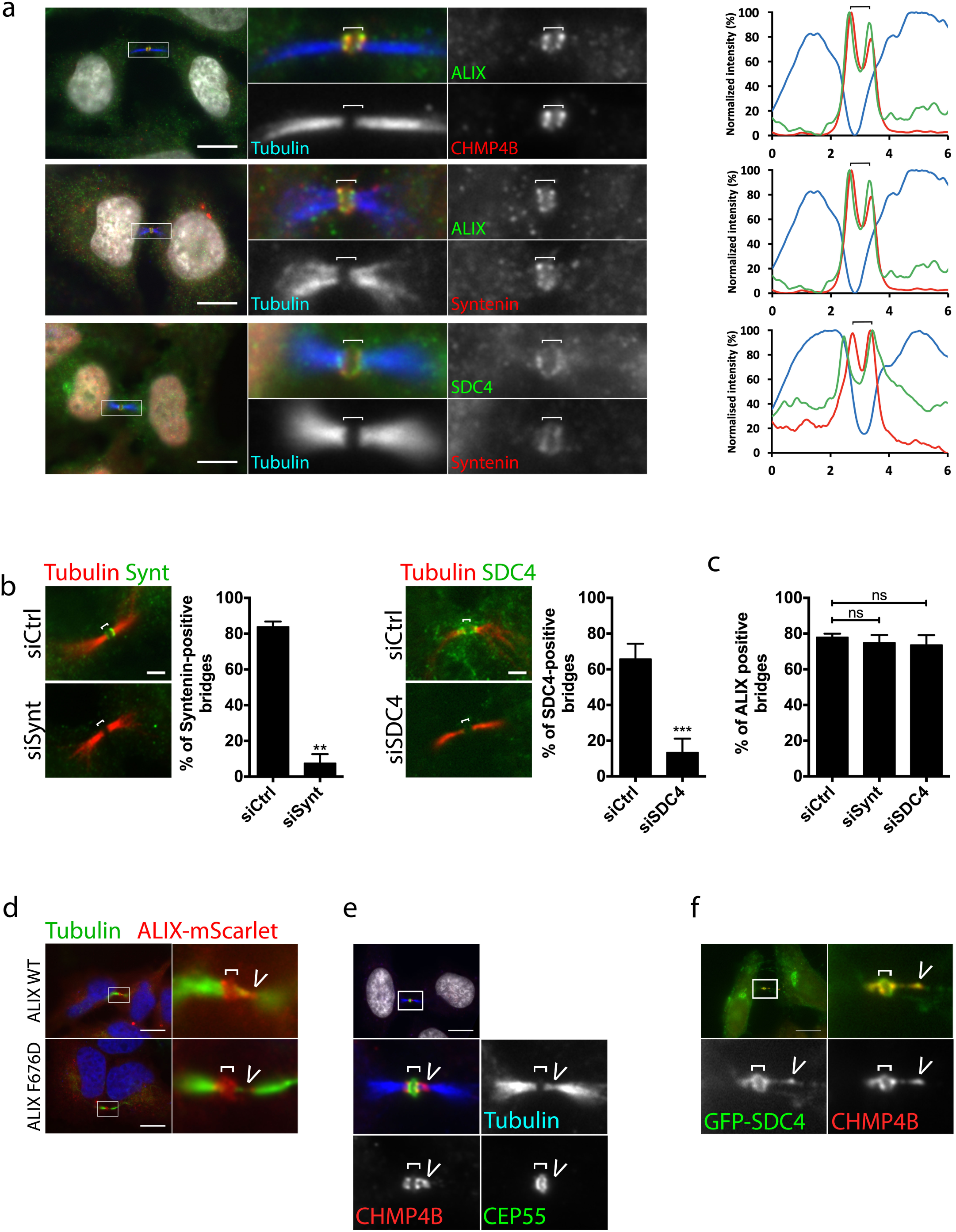
Localization of syntenin, syndecan-4 and ALIX in wild-type and depleted cells. (a) Endogenous localization of ALIX, CHMP4B, syntenin, syndecan-4 (SDC4) and acetyl-tubulin in late bridges without abscission site in fixed HeLa cells, as indicated. We used an antibody recognizing the ectodomain of SDC4 for the immunofluorescence. *Right panels:* Intensity profiles along the bridge of the corresponding images with matched colours from left panels. (b) Specificity of antibody staining. *Left panels:* late cytokinetic bridges stained for endogenous syntenin and acetyl tubulin, in control and syntenin siRNAs-treated cells, as indicated. Quantification of the % of bridges positive for syntenin in each condition. n=31-53 cells, N=3. *Right panels*: late cytokinetic bridges stained for endogenous syndecan-4 (SDC4) and acetyl tubulin, in control and syndecan-4 siRNAs-treated cells, as indicated. Quantification of the % of bridges positive for syntenin in each condition. n=22-37 cells, N=3. (c) Quantification of the proportion of bridges positive for ALIX in control, syntenin and syndecan-4 siRNAs-treated cells. n=25-64 cells, N=4. (d) Late cytokinetic bridges in HeLa cells transfected with either wild-type or F676D mutant ALIX tagged with mScarlet. Note that both construct localize to the intercellular bridge. (e) Localization of endogenous CHMP4B and CEP55 in cytokinetic bridges with abscission site, as indicated. Note that CEP55 is absent from the abscission site, where CHMP4B is localized. a, d and e: Scale bars = 10 μm. B: Scale bars = 2 μm ns: non significant, ** p < 0.01, *** p < 0.001 Student-t tests. Brackets and arrowheads mark the midbody and the abscission site, respectively. (f) Localization of endogenous CHMP4B in GFP-syndecan-4 stable cell line. The CHMP4B staining indicates that this zoomed region corresponds to a cytokinetic bridge fixed shortly before abscission. Note the colocalization between CHMP4B and syndecan-4. Bracket and arrowhead mark the midbody and the abscission site, respectively. Scale bar = 10 μm.

**Supplementary Table 1:** Proteomic and statistical analysis (Excel file). Supplementary Table 1 provides several tabs:

TAB1 The Flemmingsome or proteome of Midbody Remnants

TAB2 Enriched Flemmingsome

TAB3 Total proteins identified by mass spectrometry

TAB4 Proteins only quantified in MB+

TAB5 iBAQ values of all the proteins identified by Mass Spectrometry

TAB6 Analysis with data from the proteome from Ahna Skop (Science 2004)

**Supplementary Table 2:**
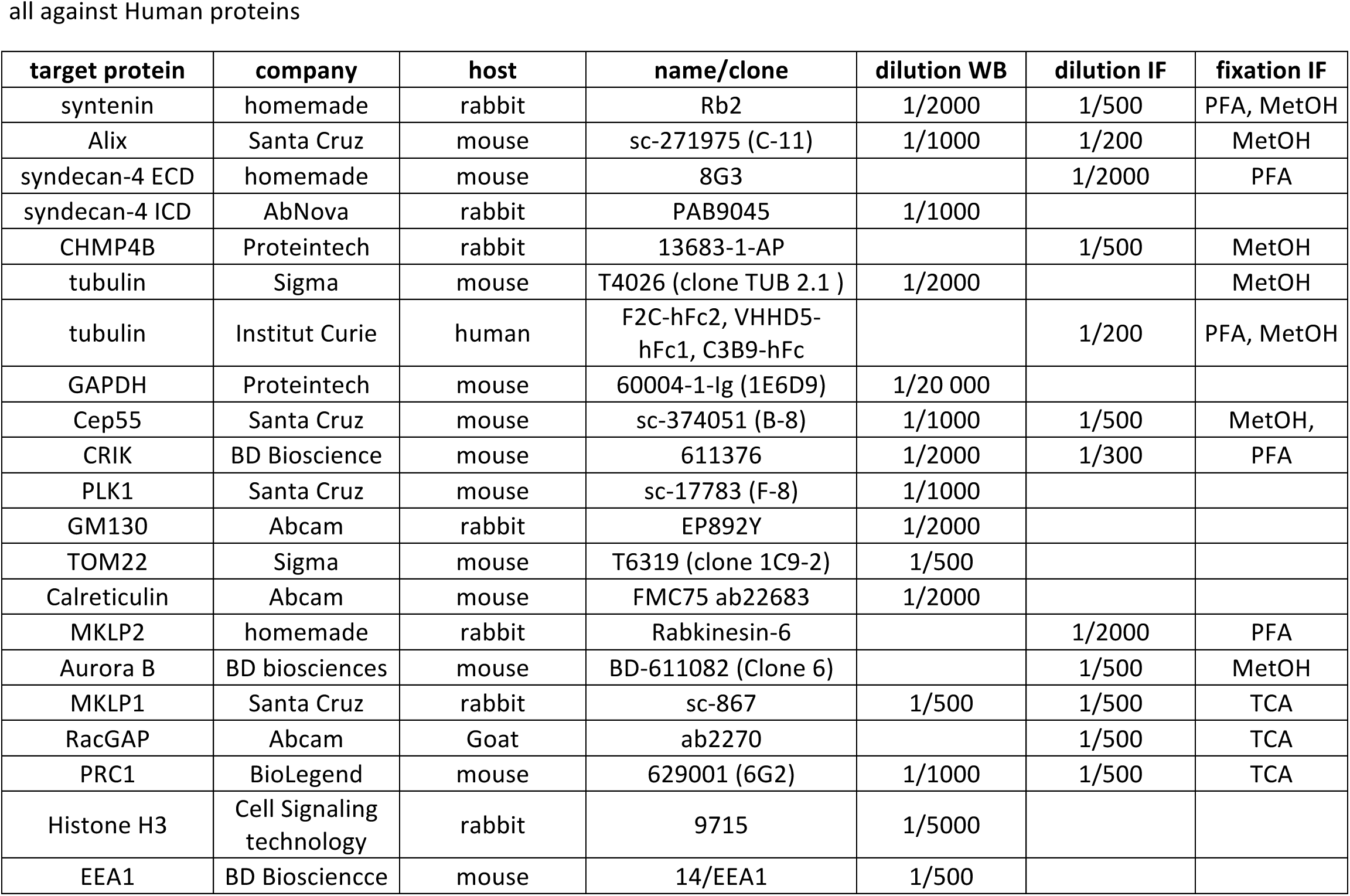
Antibodies, dilutions and fixations used in this study.

| target protein | company | host | name/clone | dilution WB | dilution IF | fixation IF |
| --- | --- | --- | --- | --- | --- | --- |
| syntenin | homemade | rabbit | Rb2 | 1/2000 | 1/500 | PFA, MetOH |
| Alix | Santa Cruz | mouse | sc-271975 (C-11) | 1/1000 | 1/200 | MetOH |
| syndecan-4 ECD | homemade | mouse | 8G3 |  | 1/2000 | PFA |
| syndecan-4 ICD | AbNova | rabbit | PAB9045 | 1/1000 |  |  |
| CHMP4B | Proteintech | rabbit | 13683-1-AP |  | 1/500 | MetOH |
| tubulin | Sigma | mouse | T4026 (clone TUB 2.1 ) | 1/2000 |  | MetOH |
| tubulin | Institut Curie | human | F2C-hFc2, VHHD5-hFc1, C3B9-hFc |  | 1/200 | PFA, MetOH |
| GAPDH | Proteintech | mouse | 60004-1-Ig (1E6D9) | 1/20 000 |  |  |
| Cep55 | Santa Cruz | mouse | sc-374051 (B-8) | 1/1000 | 1/500 | MetOH, |
| CRIK | BD Bioscience | mouse | 611376 | 1/2000 | 1/300 | PFA |
| PLK1 | Santa Cruz | mouse | sc-17783 (F-8) | 1/1000 |  |  |
| GM130 | Abcam | rabbit | EP892Y | 1/2000 |  |  |
| TOM22 | Sigma | mouse | T6319 (clone 1C9-2) | 1/500 |  |  |
| Calreticulin | Abcam | mouse | FMC75 ab22683 | 1/2000 |  |  |
| MKLP2 | homemade | rabbit | Rabkinesin-6 |  | 1/2000 | PFA |
| Aurora B | BD biosciences | mouse | BD-611082 (Clone 6) |  | 1/500 | MetOH |
| MKLP1 | Santa Cruz | rabbit | sc-867 | 1/500 | 1/500 | TCA |
| RacGAP | Abcam | Goat | ab2270 |  | 1/500 | TCA |
| PRC1 | BioLegend | mouse | 629001 (6G2) | 1/1000 | 1/500 | TCA |
| Histone H3 | Cell Signaling technology | rabbit | 9715 | 1/5000 |  |  |
| EEA1 | BD Biosciencce | mouse | 14/EEA1 | 1/500 |  |  |

**Supplementary video 1:** Colocalization of CHMP4B-GFP and ALIX-mScarlet at the midbody and at the abscission site.

HeLa cells transiently co-transfected with plasmids encoding CHMP4B-GFP and ALIX-mScarlet were recorded by spinning-disk confocal microscopy every 10 min. Merge and individual channels (in grey levels) are provided. Time 0 corresponds to the time frame preceding furrow ingression. Scale bar= 10 μm.

**Supplementary video 2:** Colocalization of GFP-syntenin and ALIX-mScarlet at the midbody and at the abscission site.

HeLa cells transiently co-transfected with plasmids encoding GFP-sytenin and ALIX-mScarlet were recorded by spinning-disk confocal microscopy every 10 min. Merge and individual channels (in grey levels) are provided. Time 0 corresponds to the time frame preceding furrow ingression. Scale bar= 10 μm.

**Supplementary video 3:** Colocalization of GFP-SDC4 and mScarlet-syntenin at the midbody and at the abscission site.

HeLa cells transiently co-transfected with plasmids encoding GFP-SDC4 and mScarlet-syntenin were recorded by spinning-disk confocal microscopy every 10 min. Merge and individual channels (in grey levels) are provided. Time 0 corresponds to the time frame preceding furrow ingression. Scale bar= 10 μm.

**Supplementary video 4:** CHMP4B-GFP behavior during cytokinesis in control cells. HeLa cells that stably expressed CHMP4B-GFP were treated with control siRNAs and recorded by spinning-disk confocal microscopy every 10 min. Time 0 corresponds to the frame preceding the arrival of CHMP4B at the midbody. The arrow indicates that abscission has occurred.

**Supplementary video 5:** CHMP4B-GFP behavior during cytokinesis in ALIX-depleted cells.

HeLa cells that stably expressed CHMP4B-GFP were treated with ALIX siRNAs and recorded by spinning-disk confocal microscopy every 10 min. Time 0 corresponds to the frame preceding the arrival of CHMP4B at the midbody. The arrow indicates that abscission has occurred.

**Supplementary video 6:** CHMP4B-GFP behavior during cytokinesis in syntenin-depleted cells.

HeLa cells that stably expressed CHMP4B-GFP were treated with syntenin siRNAs and recorded by spinning-disk confocal microscopy every 10 min. Time 0 corresponds to the frame preceding the arrival of CHMP4B at the midbody. The arrow indicates that abscission has occurred.

**Supplementary video 7:** CHMP4B-GFP behavior during cytokinesis in syndecan-4-depleted cells.

HeLa cells that stably expressed CHMP4B-GFP were treated with syndecan-4 siRNAs and recorded by spinning-disk confocal microscopy every 10 min. Time 0 corresponds to the frame preceding the arrival of CHMP4B at the midbody. The arrow indicates that abscission has occurred.

